# Shared neurogenesis onset is sufficient to explain bilateral matching in the vertebrate retina

**DOI:** 10.64898/2026.09.02.748850

**Authors:** Diana García-Morales, Adolfo Alsina, Guillaume Salbreux, Caren Norden

## Abstract

Bilateral symmetry is a hallmark of many paired organs and often essential for optimal functionality. The vertebrate eyes are a prominent example of this, as the matched development of the two retinas is required for accurate visual processing.

While macroscopic aspects of symmetry emergence across systems have been investigated, how bilateral matching is maintained once cells start to differentiate remains less understood. Here we address this question using the zebrafish retina as a model to follow neurogenic programs *in vivo* at single-cell resolution. We perform quantitative 3D live imaging of both retinas simultaneously and directly compare neurogenesis onset and propagation within and across embryos. We find that neurogenic waves initiate at the retinal poles and progress towards the mid-retina in a conserved spatiotemporal pattern. Within embryos, the two eyes exhibit highly similar neurogenesis dynamics when it comes to timing of neurogenesis onset, cell number increase, and spatial wave progression. Across embryos, however, variability is larger. While these observations hint at active inter-retinal coordination, a stochastic model predicts that a shared onset of neurogenesis can be sufficient to explain bilateral matching. Targeted genetic perturbation experiments support this prediction. We find that altering wave propagation affects patterning but not bilateral similarity. Disrupting neurogenesis onset timing, however, reduces bilateral symmetry between eyes.

Thus, the combination of experiment and theory identifies synchronized neurogenesis onset as a key determinant of bilateral symmetry, revealing a minimal principle for how reproducible development of paired organs can emerge from stochastic processes.

## Introduction

Bilaterality is a defining feature of invertebrate and vertebrate body plans. While some paired body parts like lungs, kidneys and breasts tolerate asymmetries, others need to be symmetric for optimal functionality (1–7). This is true for legs and wings where symmetry is needed for coordinated movements (1, 8–11) as well as for eyes or ears where symmetry assures optimal flow of sensory information (2, 12, 13). Currently, how bilateral matching is maintained as paired organs undergo differentiation and patterning is not fully understood.

Studies in different model organisms, notably in the Drosophila wing discs (8–11), zebrafish otic vesicle (14) and zebrafish somites have investigated bilaterality across development (15). So far, however, most of these studies concentrated on macroscopic aspects of bilateral symmetry while the question of how these macroscopic features arise from cellular patterning and fate decisions stays largely unexplored.

The paired vertebrate eyes, and particularly their light sensing neural tissues, the retinas, are excellent models to investigate these questions as they require bilateral symmetry for optimal functionality. This means that failures in developing and maintaining bilateral eye symmetry can have direct consequences for visual processing and thereby for the organism’s interactions with its environment (13, 16). The composition of the vertebrate retina is conserved over evolution and consists of five neuronal subtypes that are arranged into three distinct layers: Retinal Ganglion Cells (RGCs) make up the basal GC layer and send out the optic nerve to the tectum, Photoreceptors (PRs) reside in the apical Outer Nuclear Layer, and interneurons, Amacrine Cells, Horizontal Cells (HCs) and Bipolar Cells (BCs), in the central Inner Nuclear Layer (Fig. 1A) (17–21). These neurons emerge from multipotent progenitors during two neurogenic waves that need to generate the correct neurons at the right time in the correct proportions. During the first neurogenic wave progenitors start expressing Ath5, a transcription factor of the BHLH family (22–25). In fish and other vertebrates, these Ath5 positive progenitors give rise to RGCs, ACs, HCs and PRs (Fig. 1B) (17, 26). Later in development, but coinciding with the first, the second neurogenic wave starts, marked by the appearance of cells expressing Vsx1, a homeobox transcription factor (27, 28). Vsx1 positive progenitors mainly give rise to BCs and some ACs (Fig. 1B) (28, 29). Currently, it is not fully understood how this patterning advances and whether it is coordinated across the two eyes of the organism.

**Figure 1.**
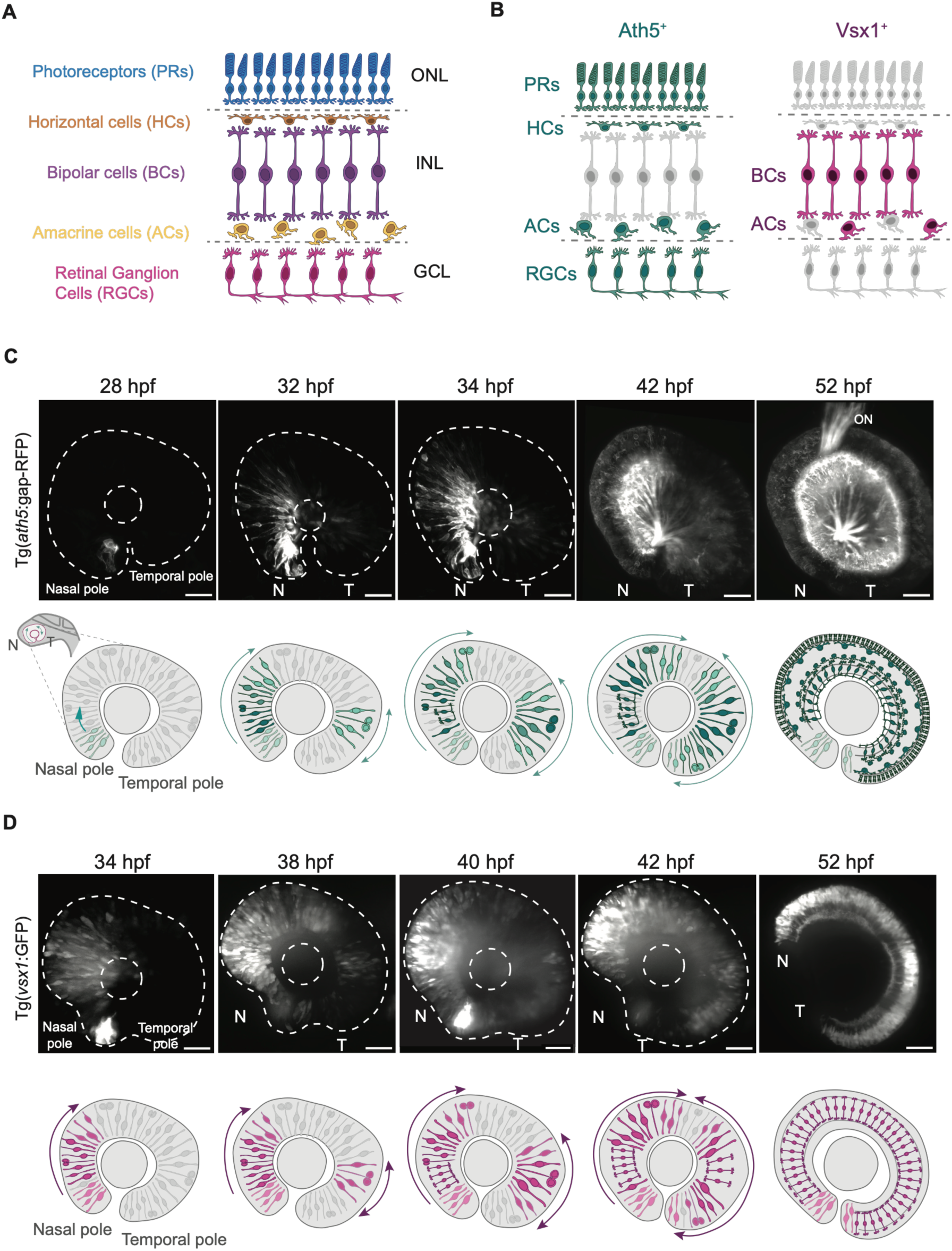
Neurogenic onset in the zebrafish retina proceeds from nasal and temporal poles toward the mid-retina. (A) Schematic of the mature zebrafish retina, showing the five neuronal cell types organized in three nuclear layers. Photoreceptors (PRs) in the outer nuclear layer (ONL), Bipolar Cells (BCs), Amacrine Cells (ACs) and Horizontal Cells (HCs) in the inner nuclear layer (INL), and Retinal Ganglion Cells (RGCs) in the ganglion cell layer (GCL). (B) Schematics illustrating the lineage output of each neurogenic wave. The Ath5 lineage (teal) produces PRs, HCs, ACs and RGCs. The Vsx1 lineage (magenta) gives rise to BCs and a subset of ACs. Greyed-out cell types indicate neurons not produced by the respective wave. (C) Representative snapshots from an *in vivo* light-sheet movie showing the onset and spatiotemporal propagation of the first neurogenic wave labelled by Tg(*ath5*:gap-RFP), between 28 hpf and 52 hpf. Ath5 expression initiates at the nasal pole (N) at 28 hpf and at 32 hpf at the temporal pole (T). Wavefronts propagate towards the mid-retina. Schematics illustrate wave progression within the retina, with undifferentiated progenitors shown in grey. ON, optic nerve. Dashed lines indicate the retinal and lens outlines. (D) Representative snapshots from an *in vivo* light-sheet movie showing the onset and propagation of the second neurogenic wave, labelled by Tg(*vsx1*:GFP) between 34 hpf and 52 hpf. Vsx1 expression initiates at the nasal pole (N) at 34 hpf and subsequently at the temporal pole (T) at 38 hpf. Wavefronts propagate towards the mid-retina. Schematics illustrate the Vsx1 wave progression within the retina. Scale bars: 20 µm. Developmental stage in hours post-fertilisation (hpf).

To get insights into these questions, the zebrafish is an ideal system, as its rapid development, transparency and small size provide the unique possibility to perform fixed and live imaging in the intact organism.

These advantages have been used in previous studies that investigated neurogenesis onset and propagation in single eyes. Here, it was postulated that neurogenesis starts at nasal positions and proceeds in a fan-like manner towards the temporal pole (20, 23, 24, 30, 31). This propagation has been linked to intra-retinal Shh signalling, where newly differentiated neurons, namely RGCs and ACs, produce Shh that promotes further neurogenic commitment in the tissue (24, 31–36). However, so far, most of these studies used fixed analysis or live imaging at low temporal resolution (23, 24, 28, 33–35). This meant that the exact dynamics and kinetics of each wave and how comparable they are within and between embryos remained unresolved.

In another line of research, lineages and fate decisions have been studied extensively (26, 37–43). In this context, previous studies revealed a degree of stochasticity in individual progenitor outcomes that assures that tissue architecture is preserved (26, 39, 40). It remains unclear, however, whether neurogenic fate decisions and ensuing lamination occur in parallel between the two eyes of an embryo or whether each eye follows its own independent programme.

In this study, we address these gaps building on previous knowledge and extend this by performing live imaging of both retinas simultaneously at single-cell resolution. This is a prerequisite for a direct quantitative comparison within and between embryos. Quantitative live imaging was combined with theory to model stochastic differentiation dynamics.

We find that both neurogenic waves follow a conserved spatiotemporal pattern across embryos. Interestingly, the two eyes within embryos are more closely matched to each other than eyes between embryos. A quantitative 3D analysis of the Ath5-positive first neurogenic wave further revealed that neurogenic cell numbers as well as spatial propagation of neurogenic cell emergence is very similar between eyes of the same embryo, however minor left-right variability remains.

Based on this quantitative analysis, our stochastic model predicted that simultaneous neurogenesis onset in both eyes rather than active inter-retinal communication, could be sufficient to account for the observed bilateral matching. This means that wave propagation thereafter can in principle proceed independently in each retina. This prediction was confirmed by experiments interfering with intraretinal Shh signalling through targeted cell depletion. Taken together, experiment and theory suggest that synchronous bilateral organ development can arise without the requirement for active inter-retinal coordination, a concept possibly relevant for other examples of paired organ development.

## Results

### Neurogenesis in the zebrafish retina proceeds from nasal and temporal poles towards the mid-retina

To reveal the dynamics of neurogenesis in the zebrafish retina over time, we imaged Tg(*ath5*:gap-RFP) embryos labelling the first neurogenic wave and Tg(*vsx1*:GFP) embryos labelling the second neurogenic wave (28, 44) individually and in combination (Fig. 1C-D and Movie S1). Light-sheet microscopy was used to minimise phototoxicity and embryos were imaged at 5 min intervals between 28 hpf, corresponding to the onset of Ath5 expression and 52 hpf, when most progenitors of first and second wave have entered neurogenesis (20, 30, 45).

We found that Ath5 positive cells first emerge at the nasal pole around 28 hpf, as previously reported (20, 23, 24, 26, 30, 31). At approximately 32 hpf, Ath5 positive cells started to also appear at the temporal pole (Fig. 1C and Movie S1). From 42 hpf the nasal and temporal wave fronts converged towards the mid-retina (Fig. 1C and Movie S1). A similar pattern was observed for the Vsx1 positive second wave, with cells appearing at the nasal pole around 34 hpf, the temporal pole from 38 hpf, and wavefronts meeting at the mid-retina at later developmental stages (Fig. 1D and Movie S1).

Our finding that neurogenesis starts at both poles and wavefronts, then merges in the mid-retina, is different to previous studies that reported a unidirectional naso-temporal wave propagation (20, 23, 24, 28, 30, 31, 46) (see discussion).

### A pipeline for quantitative analysis of neurogenic wave dynamics within and across embryos

The light-sheet protocol outlined above allowed us to simultaneously image neurogenesis onset and propagation in both eyes of a single embryo. When we qualitatively compared the onset of neurogenesis between the two eyes of the same embryo, we noted that the emergence of Ath5 as well as of Vsx1 positive cells still occurred at similar time points in each eye.

To more carefully examine bilateral neurogenesis onset and propagation, we generated two-dimensional representations of wave dynamics (Fig. 2A-A’’). To this end, we created maximum-intensity projections that were straightened along the nasotemporal axis. From these we assembled successive time points between 28 hpf and 52 hpf at 5 min intervals into a single kymograph (Fig. S1A-C). This allowed us to directly compare the spatiotemporal progression of neurogenesis between the two eyes of a single embryo as well as between embryos.

**Figure 2.**
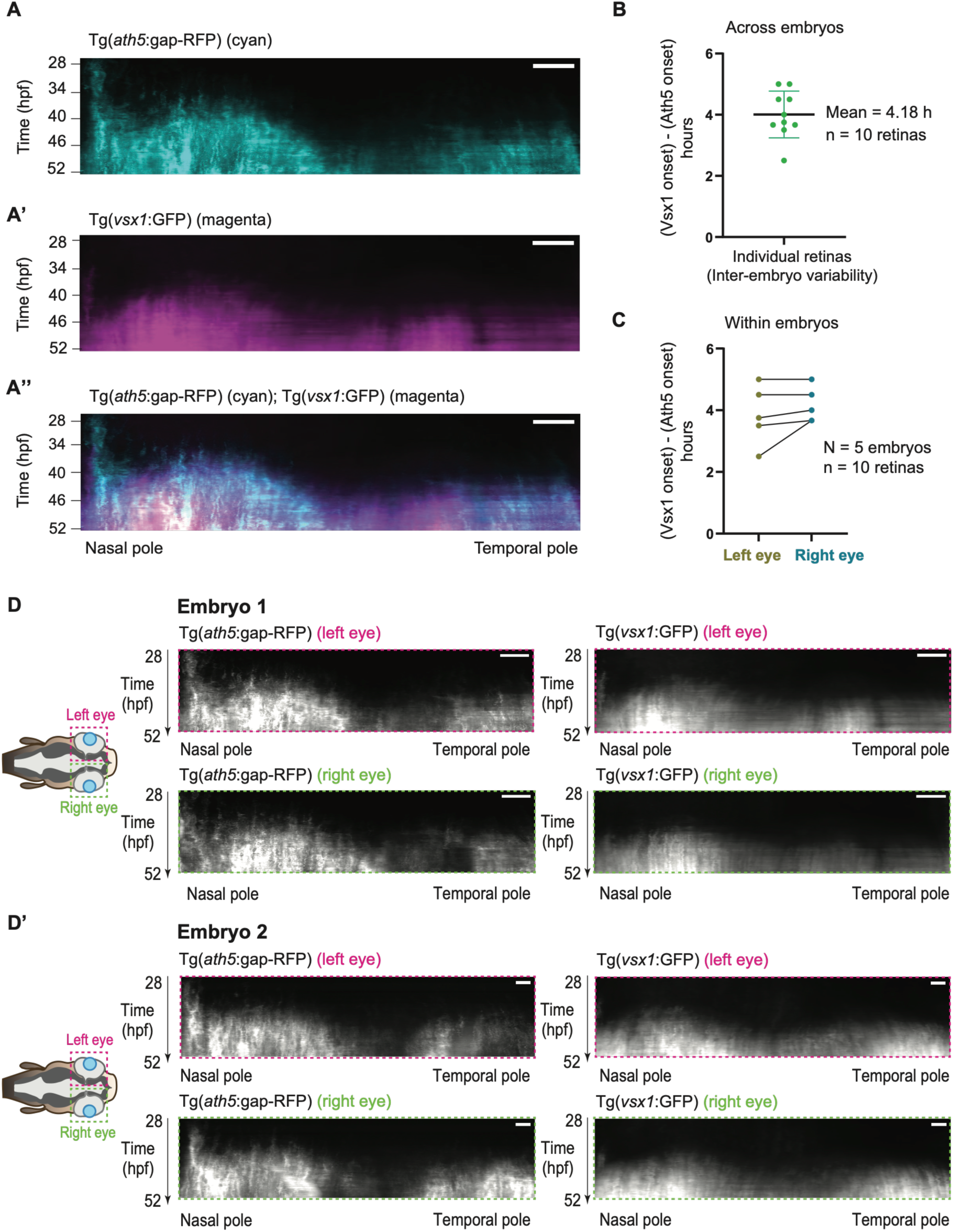
Neurogenic waves are more similar within than between embryos. (A–A’’) Representative 2D kymographs of the first and second neurogenic wave in the zebrafish retina, from 28 hpf to 52 hpf. (see Fig. S1 for details on kymograph generation) (A) Spatiotemporal propagation of the Ath5 wave, labelled by Tg(*ath5*:gap-RFP). (A’) Spatiotemporal propagation of the Vsx1 wave, labelled by Tg(*vsx1*:GFP). (A’’) Overlay of both channels showing the sequential expression of Ath5 (cyan) and Vsx1 (magenta). In all kymographs, the x-axis represents the position along the naso-temporal axis, and the y-axis represents developmental stage (28 hpf – 52 hpf). (B) Delay between Ath5 and Vsx1 wave onset across retinas (inter-embryo variability). Each dot represents one retina; mean delay is 4.18 h (n = 10 retinas from N = 5 embryos). (C) Paired comparison of the Ath5-to-Vsx1 onset between left (olive) and right (turquoise) eyes within individual embryos. Lines connect the two eyes of the same embryo (N = 5 embryos, n = 10 retinas). (D–D’) Representative kymographs from two different embryos showing Ath5 (left column) and Vsx1 (right column) spatiotemporal dynamics in the left (top) and right (bottom) eyes. Propagation dynamics are qualitatively more similar within embryos than between embryos. Scale bars: 20 µm. Developmental stage in hours post-fertilisation (hpf).

These kymographs confirmed that both neurogenic waves propagated from the nasal and later temporal poles toward the mid-retina (Fig. 2A-A’’, D-D’). Ath5 positive cells started to emerge at around 28 hpf, followed by the Vsx1 positive cells at around 34 hpf with an average 4 h delay between waves (Mean = 4.18h, SD = 0.59h) (Fig. 2B; N = 5 embryos, n = 10 retinas). The delay between Ath5 and Vsx1 onset roughly matched between the left and right eyes of the same embryo (Fig. 2C; N = 5 embryos, n = 10 retinas) with lower variability than between embryos (Fig. 2B, C). Further, qualitative comparison of kymographs across waves and embryos showed that the patterns of wave onset and propagation were more similar within than between embryos (Fig. 2D-D’).

To determine whether this qualitative similarity was reflected by similar numbers of neurogenic cells emerging in the two eyes of the same embryo, we counted Ath5 positive cells at 30 hpf in fixed whole-mount retinas across the sections of a full z-stack (Fig. S2; N= 17 embryos; n = 34 retinas). We focused on the first neurogenic wave, as it gives rise to the majority of retinal neurons (RGCs, ACs, HCs, and PRs) (Fig. 1B) (26, 40, 42, 47, 48), while still presenting low cell density at 30 hpf (Fig. S2A).

These quantifications revealed that cell numbers were more similar between the two eyes of the same embryo than between embryos. However, the variability of emerging cells spanned a wide range across embryos from 1 to 212 cells at 30 hpf (Fig. S2B). Due to the limitations of fixed samples, we could not differentiate whether these differences arose because the embryos were not all at the same developmental stage at fixation or because neurogenesis initiates within a broader time window. At later stages from 34 hpf our counting approach was no longer feasible as cells could not be unambiguously differentiated due to the use of the Tg(*ath5*:gap-RFP) membrane marker (Fig. S2C, D).

To overcome these constraints and better quantify neurogenesis onset and propagation within and across embryos, we generated a nuclear Ath5 reporter line Tg(*ath5*:H2B-mNeonGreen) using an established Ath5 promoter sequence (see Methods) (47, 49). Nuclear signals provide spatially discrete and individually resolvable reference points and thus allow for the segmentation of cells in increasingly crowded tissues (50, 51).

Initial characterisation of this transgenic line confirmed that the nuclear Ath5 reporter expression overlaps with cells labelled by the Tg(*ath5*:gap-RFP) membrane marker (Fig. 3A).

**Figure 3.**
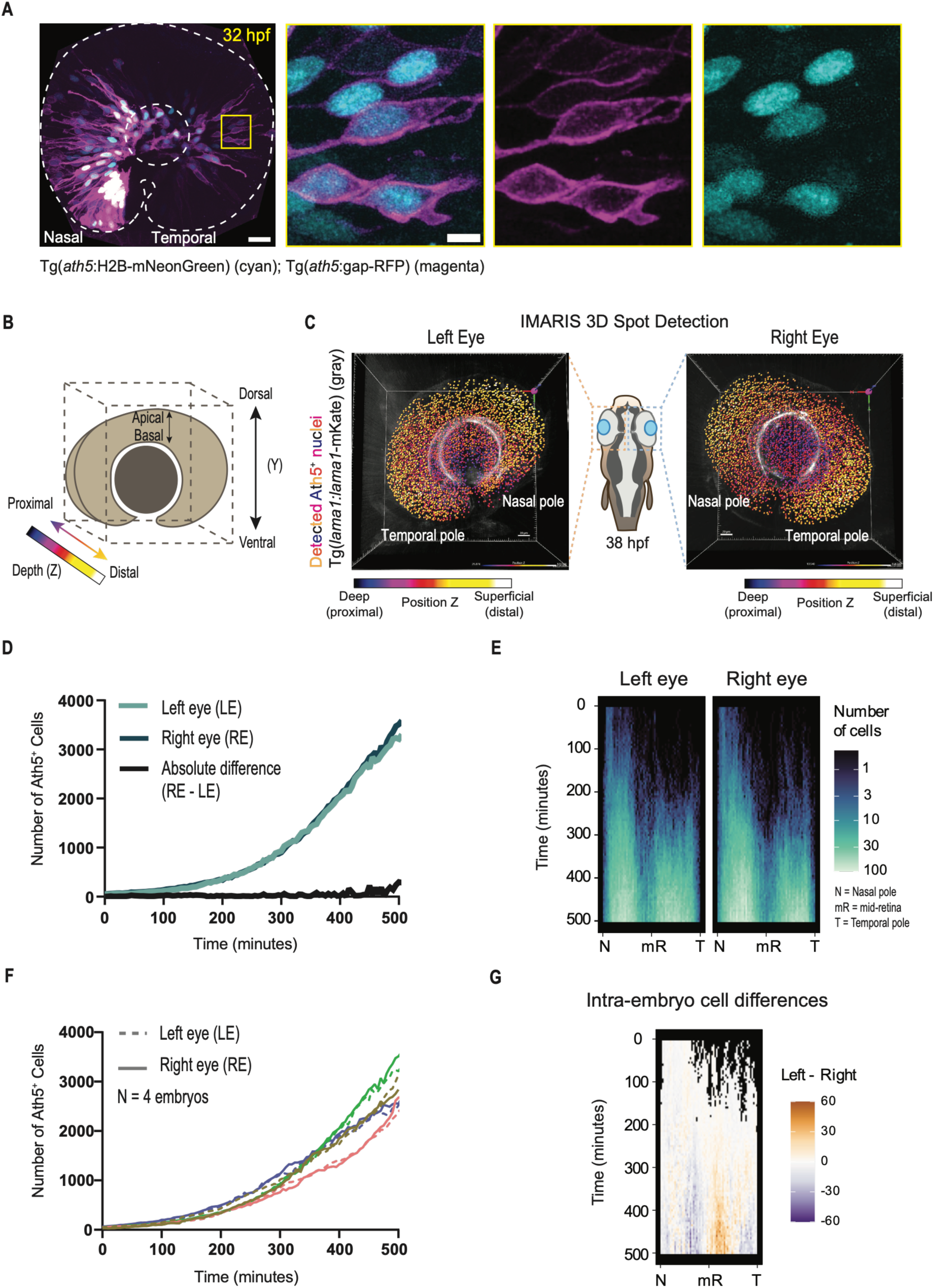
Neurogenic cell emergence and wave propagation are similar in the two eyes of the same embryo. (A) Confocal image of a zebrafish retina at 32 hpf showing Ath5^+^ cells labelled by Tg(*ath5*:gap-RFP) (magenta, membranes) and Tg(*ath5*:H2B-mNeonGreen) (cyan, nuclei). The yellow inset shows a close-up of the boxed region, with the merged signal of the individual channels. (B) Schematic of the 3D coordinates used for depth colour-coding in (C). The y-axis represents the dorsal-ventral axis (top to bottom), the x-axis represents the naso-temporal axis (right to left), and the z-axis represents the proximal-to-distal axis (deepest to surface). (C) 3D reconstruction of paired left and right retinas at 38 hpf, showing the retinal outline labelled by Tg(*lama1:lama1*-mKate2) (grey) and the detected Ath5^+^ nuclei colour-coded by depth along the z-axis (FIRELUT: proximal = purple/cool colours, deepest region; distal = yellow/warm colours, superficial region). (D) Number of Ath5^+^ cells over time in the left (light teal) and right (dark teal) eyes of a representative embryo. Spot detection and quantification were performed for 500 minutes after the first eye reached 50 Ath5^+^ cells. The black line shows the absolute difference between the left and right eyes (RE - LE) at each time point. (E) Spatiotemporal kymographs showing the number and distribution of Ath5^+^ emergent cells generated from the 3D single-nucleus detection (x-axis; N, nasal pole; mR, mid-retina; T, temporal pole) (y-axis; 0-500 minutes; 28-36.3 hpf) for the left and right eyes of the embryo shown in (D). The number of Ath5^+^ cells is colour-coded ranging from 0 (black) to 100 (white). (F) Number of Ath5^+^ cells over time in left (dashed) and right (solid) eyes for N = 4 embryos. (G) Difference kymograph between the left and right eyes along the naso-temporal axis over time for the embryo shown in (D, E). Differences are colour-coded from - 60 (purple, more cells in the right eye) to 60 (orange/brown, more cells in the left eye), with white indicating no difference and black corresponding to positions where no cells were detected in either eye. Scale bar: 20 µm (A, C). Time in panels D-G in minutes.

To limit our analysis to cells within the retina, we combined the Tg(*ath5*:H2B-mNeonGreen) line with Tg(*lama1:lama1*-mKate2), which labels the extracellular matrix surrounding the emerging retina (52, 53) (Fig. 3C and Movie S2). Light-sheet imaging between 28 hpf and 36 hpf at 5 min intervals was used to generate high-resolution 4D (XYZT) datasets of the first neurogenic wave (Movie S2).

To reliably detect Ath5 positive nuclei in these datasets, we initially tested deep-learning-based approaches, including our previously established StarDist protocol (51). However, the lower axial resolution of light-sheet microscopy combined with variable nuclear signal intensity led to systematic detection errors, including missed and merged nuclei. We also tested CellPose, but once more the heterogeneous nuclear signal in our datasets caused unreliable segmentation across both time and z-depth.

To overcome these limitations, we developed a custom-made 3D analysis pipeline based on IMARIS spot detection (Fig. 3B, C and Movie S2) that identified the centroid of each nucleus filtered by the mean fluorescence intensity. This method was substantially more accurate than the approaches described above, particularly during early neurogenesis (from 28 hpf - 32 hpf). After 32 hpf, we introduced a manual curation step (see Methods).

The IMARIS pipeline now allowed for a quantitative comparison of neurogenic output (Fig. 3D and Fig. S3A), timing (Fig. S3B) and spatial propagation (Fig. 3E and Fig. S3C) of the first neurogenic wave within and across embryos.

### Neurogenesis onset and propagation of the first Ath5 positive wave are more similar within than between embryos

Using the IMARIS platform we quantified emerging Ath5 positive cells in each eye over the first 500 min after neurogenesis onset starting at 28 hpf. To normalize developmental stage across embryos, time 0 was defined as the point at which the first eye of the pair reached 50 Ath5-expressing cells (N = 4 embryos).

After 500 min, eyes contained between 2400 and 3800 Ath5 positive cells (Fig. 3F and Fig. S3A).

When comparing eyes of the same embryo, cumulative Ath5 positive cell numbers increased at nearly identical rates leading to overlapping curves during the complete imaging period (Fig. 3D and Fig. S3A). We calculated the absolute difference in Ath5 positive cell number between paired retinas at each time point (see Methods) and noted that absolute differences remained low at early stages below 50 cells for the first 300 minutes of imaging across all four embryos (Fig. 3D and Fig. S3A). Differences increased at later stages as Ath5 positive cell numbers accumulated, reaching a maximum of 299 cells at 500 minutes in the embryo with the largest bilateral difference, where the right eye contained 3510 and the left eye 3211 Ath5 positive cells respectively (Fig. 3D).

When Ath5 positive cell emergence was compared between embryos, higher variability was observed (Fig. 3F and Fig. S3A). While the overall shape of cumulative cell number curves was similar across embryos, variability increased at later stages of the imaging window (Fig. 3F).

Together, these data show that while the overall dynamics of neurogenic cell emergence are conserved across embryos, intra-individual variability was lower than inter-individual variability.

### Differentiation rates of the first neurogenic wave follow a conserved temporal programme

To further determine the dynamics of Ath5 positive cell emergence within and across embryos, we calculated their emergence rate by calculating the first derivative of the cumulative cell count (Fig. S3B). Across all embryos analysed, Ath5 positive cell emergence was similar, remaining low during the first 200 min of differentiation at less than 6 cells per min. Rates then increased progressively, reaching ranges of 7 to 13 cells per min between 200 min and 300 min. These rates continued to rise between 300 min and 380 min, reaching 7 to 17 cells per min and ranging between 11 and 27 cells per min after 400 min. When compared within embryos, differentiation rates showed closely matched dynamics throughout the imaging period (Fig. S3B).

This shows that neurogenesis during the first wave follows a conserved profile of progressively increasing differentiation rates. These rates are more similar between the two eyes of the same embryo than across embryos.

### Spatial propagation of the Ath5⁺ neurogenic wave is overall conserved and more similar within than between embryos

Having established that Ath5 positive cell numbers and differentiation kinetics match between the two eyes of the same embryo, we asked whether cells also emerge at similar spatial positions.

To analyse this, we generated spatiotemporal kymographs from the 3D single-cell detection analysis, projecting the position of individual Ath5 positive cell numbers in 2D onto the naso-temporal axis over time (see SI) (Fig. 3E and Fig. S3C).

These kymographs showed the consistent pole-to-mid-retina propagation pattern already seen in the intensity projection kymographs (Fig. 2A, Fig. 3E and Fig. S3C). When comparing the number of cells emerging across space and time, the distribution was very similar within embryos while general dynamics were conserved between embryos (Fig. S3C).

To quantify deviations of Ath5 positive cell numbers across space and time for the two eyes of the same embryo, we generated kymographs of the cell number difference (Fig. 3G and Fig. S3D). We found that differences in Ath5 positive cell numbers across space remained small throughout wave propagation during the first 300 minutes of differentiation. Furthermore, no systematic bias towards either eye or an accumulation of differences over time was observed within or across embryos.

Thus, neurogenic wave propagation follows a conserved spatiotemporal pattern across embryos. Within embryos neurogenesis propagation is closely matched for emerging cell numbers as well as spatial distribution and progression.

### A stochastic model of independent wave propagation reproduces bilateral similarity between eyes

As we showed that neurogenic onset and propagation are closely matched between eyes of the same embryo, we wondered whether this could hint at an active coordination between the two retinas or whether it would be possible for retinas that follow the same program independently.

To extract whether the variability in the differentiation process observed within a single eye is predictive of the variability between two sister eyes, we formulated a stochastic model of Ath5 positive cell emergence (Fig. 4A, B). In this model, individual cells initiate differentiation independently, with no communication between cells or between eyes (Fig. 4A). The differentiation time of each cell *τ*, measured relative to a common reference time of onset of differentiation t=0, is taken as a stochastic variable drawn from a distribution that depends on its position along the retina (see SI for details) (Fig. 4B-D). The distribution of differentiation times fully specifies the system dynamics, as the number of Ath5 positive cells at time t corresponds to the number of cells with differentiation times shorter than or equal to t (Fig. 4A). To account for the spatial dependence of the differentiation dynamics, we partitioned the retina into sections along the naso-temporal axis, each containing *N_φ_* cells. The model was then fitted to our four individual control 3D datasets (Fig. 3 and Fig. S3). In each of these embryos, the model faithfully recapitulated the number of emerging Ath5 positive cells (Fig. 4E), their rate of differentiation (Fig. 4F) and their spatial propagation from the poles towards the mid-retina (Fig. 4G). Fitting the model to experimental data resulted in a position-dependent differentiation probability that is highest at the retinal poles (Fig. 4C), consistent with the naso-temporal wave propagation seen in experiments (Fig. 3E and Fig. S3C). The average typical activation time across control embryos obtained from fitting the model is 1329 minutes, indicating that our experimental measurements span the early stages of the differentiation process.

**Figure 4.**
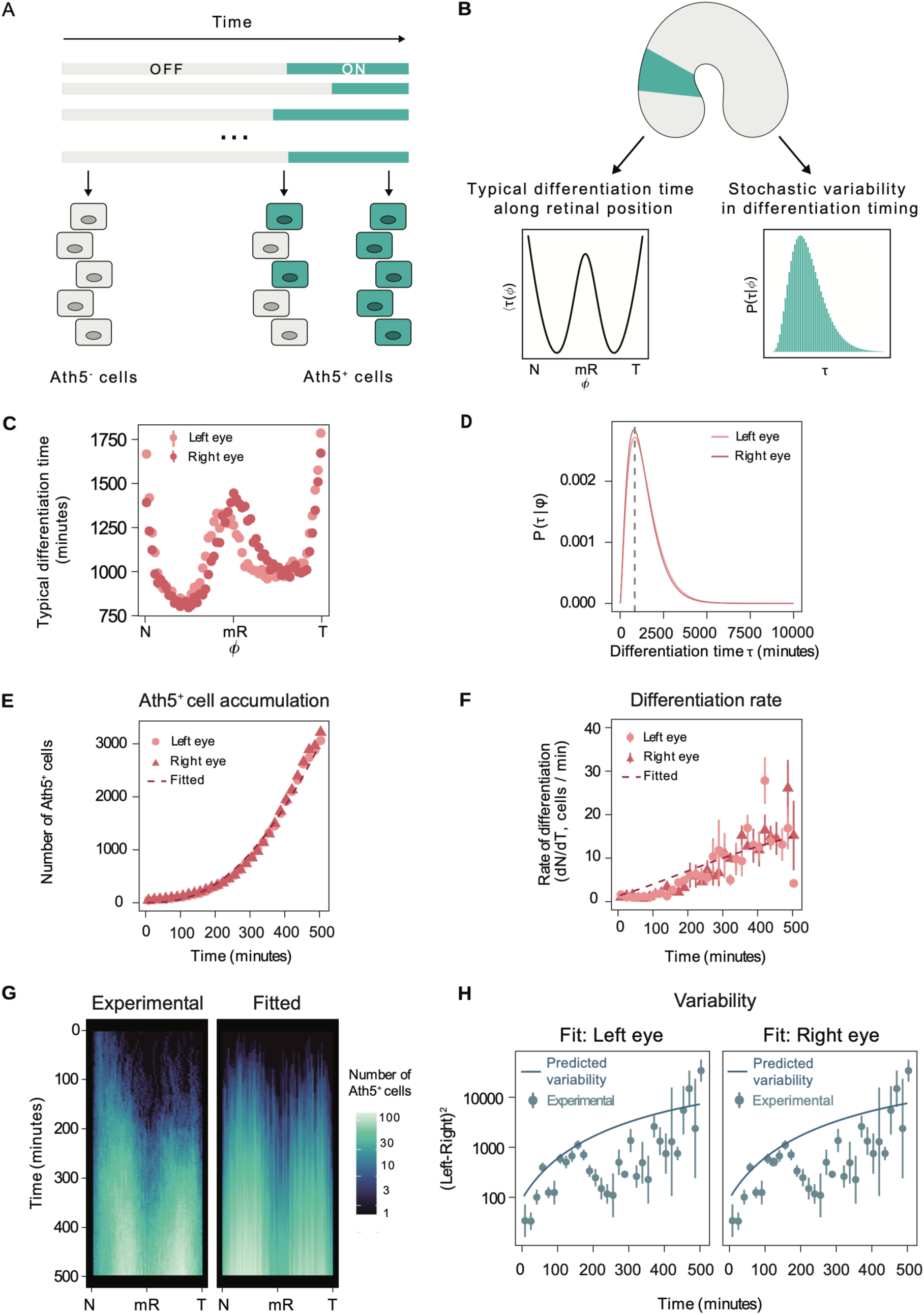
A stochastic model of cell-autonomous differentiation reproduces the dynamics and intra-embryo variability of the first neurogenic wave. (A) Schematic illustrating the stochastic differentiation process. Individual cells transition independently from an Ath5^-^ (OFF) to an Ath5^+^ (ON) state at stochastic differentiation times (τ). Once activated, Ath5 expression is maintained for the duration of the simulation. (B) Schematic representation of the theoretical model. The retina is partitioned along the naso-temporal axis (ϕ) into spatial regions, each characterised by a differentiation time distribution *P*(*τ*|*φ*). The mean differentiation time ⟨*τ*⟩(*φ*) varies along the naso-temporal axis (left), with cells at the poles having shorter mean differentiation times, consistent with earlier neurogenesis onset at these positions. The distribution *P*(*τ*|*φ*) is modelled as a Gamma distribution with parameters α**_φ_** (shape) and θ (scale) (right). (C) Typical differentiation time ⟨τ⟩(ϕ) along the naso-temporal retinal axis (ϕ), for the left (light red) and right (dark red) eyes of the representative embryo shown in (E-G). Values correspond to the average typical differentiation time for each spatial region, averaged over 100 samples from the posterior parameter distribution. Shorter typical differentiation times at the nasal (N) and temporal (T) poles reflect the earlier onset of Ath5 expression at these positions, consistent with the naso-temporal propagation of the neurogenic wave. (D) Fitted differentiation time distributions P(τ|ϕ) for the left (light red) and right (red) eyes of the same representative embryo shown for ϕ = 1.55 (E) Model-predicted Ath5^+^ cells over time compared to experimental data from paired retinas. Experimental data for left (light red circles) and right (dark red triangles) eyes of a representative embryo are shown alongside the average cell count fitted from 100 independent samples from the posterior parameter distribution for each of the two eyes (dashed line). Average parameters from the posterior distribution for the two eyes of this embryo are N_ϕ_ = 622.50, θ = 574.01 min, <α_ϕ_)=2.90 (left eye) and N_ϕ_ = 646.19, θ=552.85 min, <α_ϕ_) = 2.96 (right eye). (F) Model-fitted differentiation rate over time compared to experimental data from paired retinas. The differentiation rate is shown in cells per min for left (light red circles) and right (dark red triangles) eyes of the same representative embryo, alongside the average differentiation rate fitted from 100 independent samples from the posterior distribution of parameters for each of the two eyes (dashed line). Experimental differentiation rates were calculated from the temporal derivative of cumulative Ath5^+^ cell numbers. (G) Comparison between experimental (left) and model-fitted (right) spatiotemporal propagation dynamics of Ath5^+^ cells for the embryo shown in (C). Kymographs show the number of Ath5^+^ cells (cyan) per naso-temporal position (x-axis) over time (y-axis). Colour scale indicates the number of Ath5^+^ cells (1–100). N, nasal pole; mR, mid-retina; T, temporal pole. The model kymograph was generated from a single realisation of the stochastic process characterised by a single sample of the posterior distribution. (H) Comparison between experimentally measured intra-embryo variability and the variability predicted by the model. Experimental variability was quantified as the squared difference in Ath5^+^ cell number between left and right eyes over time (dots). The model-predicted variability (solid lines) was calculated as the expected squared difference between two independent realisations of the same stochastic process fitted to the eye indicated in the panel label (see Supplementary Theory), averaged over 100 samples from the posterior parameter distribution. Time is shown in minutes relative to the onset of differentiation (t = 0 defined as the time point at which the first eye of the pair reaches 50 Ath5^+^ cells).

Using these fitted parameters, the model allowed us to test whether independent stochastic differentiation with no communication between eyes, would be sufficient to reproduce the observed similarity and remaining variability seen between the two eyes of the same embryo. Using the fitting results from one eye, we calculated the theoretically expected variability between two independent realisations of the same stochastic process. This was then compared to the experimentally observed left-right variability in cell numbers (see SI). We found that the model accurately predicted the observed variability between sister eyes (Fig. 4H). Independent stochastic differentiation in each eye, starting from a shared onset time, can thus be sufficient to account for both the differentiation dynamics within each eye and the variability between sister eyes. In this modelling outcome, further coordination between left and right eyes would not be needed beyond a common starting point.

### Depletion of intra-retinal Shh signalling by suppressing RGC and AC emergence alters wave patterning without substantially affecting bilateral similarity of neurogenesis

As the model suggests that each retina could propagate through neurogenesis independently when starting from a shared onset condition, we asked whether intra-retinal Shh signalling, which is known to drive wave propagation within eyes (24, 31–33, 35), can influence wave dynamics and/or bilateral matching.

In controls, Shh expression showed a similar onset and propagation profile as seen for Ath5 positive neurogenic cells, with Shh signal emerging initially at the nasal (32, 33) and temporal poles before wavefronts met in mid-retinal positions (Fig. S4A). Compared to the Ath5 onset (Fig. 1C), the Shh onset was delayed (32) occurring between 36 hpf and 37 hpf (Fig. S4A).

To interfere with intra-retinal Shh we eliminated its primary sources, RGCs and ACs. To this end, we knocked down the neurogenic bHLH transcription factors Ath5 (resulting in suppression of RGC emergence) and Ptf1a (resulting in suppression of AC and HC emergence) using a previously established morpholino approach (17, 26, 54–56) (Fig. 5A). We confirmed that the morphants did not feature RGC or AC (Fig. S4B). Without these cell types present, Shh expression was reduced compared to controls (Fig. 5B). We used the Tg(*ath5*:H2B-mNeonGreen) line to monitor the emergence of neurogenic cells in RGCs/ACs-depleted embryos. This is possible as the morpholino depletes Ath5 protein but does not affect promoter activity. The Tg(*ath5*:H2B-mNeonGreen) transgene is therefore still expressed in cells in which the *ath5* promoter is active, allowing us to track cells that would normally initiate Ath5 positive neurogenesis even in the absence of Ath5 protein. Our analysis revealed altered neurogenesis dynamics compared to controls. In all RGC/AC-depleted embryos, the final number of Ath5 positive cells emerging in the first 500 min of neurogenesis was substantially reduced, ranging from approximately 800-2200 Ath5 positive cells per eye (Fig. 5C and Fig. S4C), compared to 2400-3800 in controls (Fig. 3F, Fig. 5C and Fig. S4C). Consistent with the reduction in total neurogenic output, neurogenesis rates were lower than in controls. During the first 200 min, differentiation rates rarely exceeded 4 Ath5 positive cells per min, compared with 6 cells per min in controls. After 400 min, differentiation rates ranged between 7 and 23 cells per min, compared to 11 and 27 cells per min in controls (Fig. S4F and Fig. S3B). Thus, cumulative neurogenic output was reduced, and neurogenesis progressed more slowly in the RGC/AC depletion condition.

**Figure 5.**
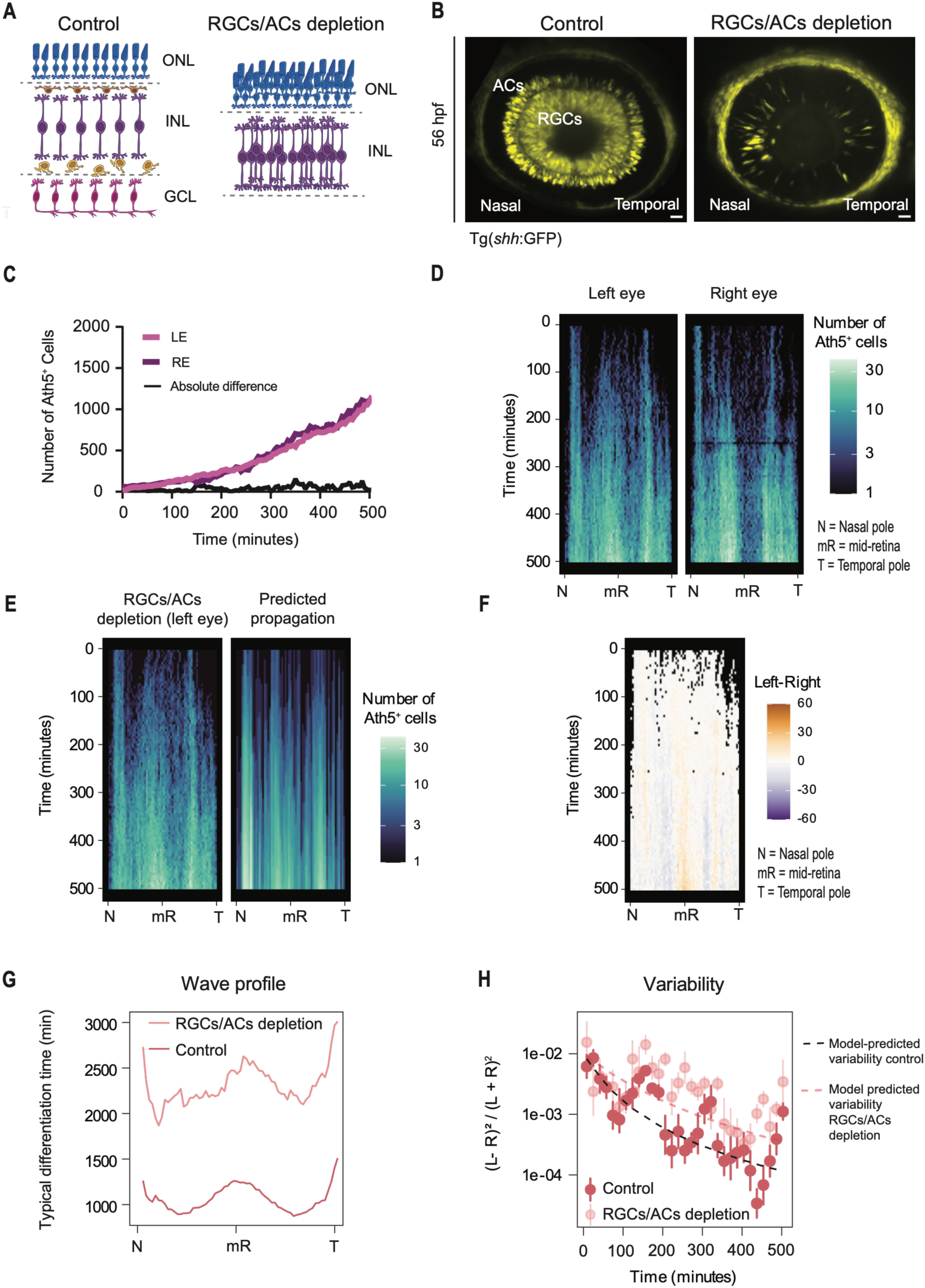
Depletion of retinal ganglion and amacrine cells alters wave propagation dynamics while generally preserving bilateral similarity of neurogenesis. (A) Schematic representation of the experimental perturbation. Left, control retinas contain all five neuronal cell types across the ganglion cell layer (GCL), inner nuclear layer (INL) and outer nuclear layer (ONL). Right, simultaneous depletion of RGCs (Ath5-MO) and interneurons (ACs and HCs; Ptf1a-MO) results in retinas lacking these cell types, containing only bipolar cells (BCs) and photoreceptors (PRs). (B) Representative confocal images of Tg(*shh*:GFP) expression at 56 hpf in a control embryo (left) and RGCs/ACs depletion condition (right). In controls, Shh is detected in RGCs (inner layer) and ACs (outer layer). In RGCs/ACs depleted embryos, Shh expression is strongly reduced, consistent with the depletion of its primary cellular sources. RGCs, retinal ganglion cells; ACs, amacrine cells. Scale bars: 20 µm. (C) Quantification of Ath5^+^ cell emergence over time in the left (light magenta) and right (dark magenta) eyes of a representative embryo in the RGCs/ACs depletion condition. The black line represents the absolute difference in the number of Ath5^+^ cells between paired eyes (RE - LE). (D) Spatiotemporal kymographs showing the number of Ath5^+^ cells along the naso-temporal axis over time in the left and right eyes of the representative RGCs/ACs-depleted embryo shown in C. Colour scale indicates cell number (1–30). N, nasal pole; mR, mid-retina; T, temporal pole. (E) Comparison of the experimental spatiotemporal kymograph from the left eye of a representative RGCs/ACs-depleted embryo (left) and the corresponding kymograph generated from the fitted model (right). The model kymograph was generated from a single realisation of the stochastic process characterised by a single sample of the posterior distribution. (F) Intra-embryo kymograph of the difference in Ath5^+^ cell number between the left and right eyes along naso-temporal positions for the depleted embryo shown in D. Differences are colour-coded from -60 (purple, more cells in the right eye) to 60 (orange, more cells in the left eye), with white indicating no difference. Black regions indicate positions and time points at which Ath5^+^ cells are absent in both eyes. (G) Typical differentiation time profile along the naso-temporal axis, as inferred by the model, for a representative RGCs/ACs-depleted embryo (light red) compared with a control embryo (dark red). The RGCs/ACs depleted condition shows a flatter profile (quantified by the standard deviation of the differentiation time profile from π/4 to 7π/4; σ_Control_ = 162.72 min, σ_RGC/AC_ = 132.13 min) and delayed differentiation times across the retina, reflecting the loss of the pole-early differentiation dynamics observed in the controls. Each profile corresponds to the average across all drawn samples of the posterior distribution for the two eyes of the corresponding embryo. (H) Quantification of the relative intra-embryo variability over time in control (dark red dots) and RGCs/ACs depletion condition (light red dots) compared to model predictions (dashed lines). Model-predicted relative variability for controls (black dashed line) and after replacing only the typical differentiation time profile with the corresponding morphant profile while keeping all other parameters fixed (red dashed line) is shown. Model predictions correspond to the average variability expected over 100 samples from the posterior parameter distribution for each eye. Time is shown in minutes from the onset of differentiation (t = 0 defined as the time point at which the first eye reaches 50 Ath5^+^ cells).

Despite this lower differentiation rate, the number of emerging Ath5 positive cells remained similar between the two eyes of the same embryo (Fig. 5C and Fig. S4C). Variability between paired eyes in the RGCs/ACs-depleted embryos was slightly increased relative to controls, with absolute differences first exceeding 50 Ath5 positive cells after approximately 110 min to 150 min of imaging (Fig. 5C, Fig. S4C). When analysing the spatiotemporal kymographs in the RGC/AC depletion condition, we noted, however, that the wave pattern was altered. Rather than initiating at the poles and propagating towards the mid-retina, Ath5 positive cells emerged across the retina more evenly (Fig. 5D and Fig. S4D). Nevertheless, spatiotemporal propagation remained closely matched between the two eyes of the same embryo albeit at much lower output levels (Fig. 5F and Fig. S4E-E’).

To test whether the modest increase of intra-embryo variability observed in RGC/AC depleted embryos could be linked to the altered spatial propagation, we asked whether a flattening of the typical differentiation time profile alone would be sufficient to increase variability in the stochastic model, when keeping all other parameters unaltered. Replacing only the mean differentiation time profile with the flatter morphant shape while keeping all other parameters at control values (see SI), the model reproduced both the altered propagation dynamics and the increase in relative variability without further adjustment (Fig. 5E, G, H). Thus, the model indicates that the increase in relative variability as observed in RGC/AC depleted retinas can be attributed solely to altered patterning of the wave.

Thus, the combination of experiment and theory indicates that RGC/AC depletion resulting in reduced intra-retinal Shh signalling influences the spatial pattern of the neurogenic wave. However, the two eyes are still closely matched in neurogenic output. The modest increase in variability observed in RGC/AC depleted embryos can be explained by the altered spatial dynamics of differentiation, without a need for additional bilateral coordination mechanisms. This suggests that spatial patterning of the wave and bilateral similarity are separable traits.

### RGC depletion increases variability of neurogenic onset between eyes by reducing bilateral symmetry while preserving propagation dynamics

So far, model and experiment suggest that wave propagation and bilateral similarity of neurogenesis are separable traits and that bilateral similarity is maintained, albeit with a small increase in variability, even when intra-retinal Shh signalling is reduced by RGC/AC depletion.

Another prediction of the model was that a shared neurogenesis onset is essential for bilateral matching, as all cells are assumed to set their differentiation time relative to a common starting point. To test this model prediction directly, we depleted Ath5 and thereby RGC emergence using an established morpholino approach (17, 26, 55, 56). We assumed that this would disrupt earliest neurogenesis propagation based on our previous finding that Ath5 knockdown shifts early neurogenic output from RGCs towards photoreceptors and inhibitory neurons, the latter, however, produced in much smaller numbers than RGCs in controls (26) (Fig. 6A). Based on the fact that RGCs and ACs are the primary sources of intraretinal Shh (Fig. S5A) (32, 34) and that after initial neurogenesis onset intraretinal Shh is required for early wave propagation (23, 31–33, 37), we expected that this strong reduction of early intraretinal Shh-producing cells would make early neurogenesis propagation more susceptible to stochastic differences between eyes. We further speculated that once sufficient neurogenic Shh-producing cells had accumulated as more ACs are produced at later developmental stages (26), propagation would proceed more similarly to the control condition.

**Figure 6.**
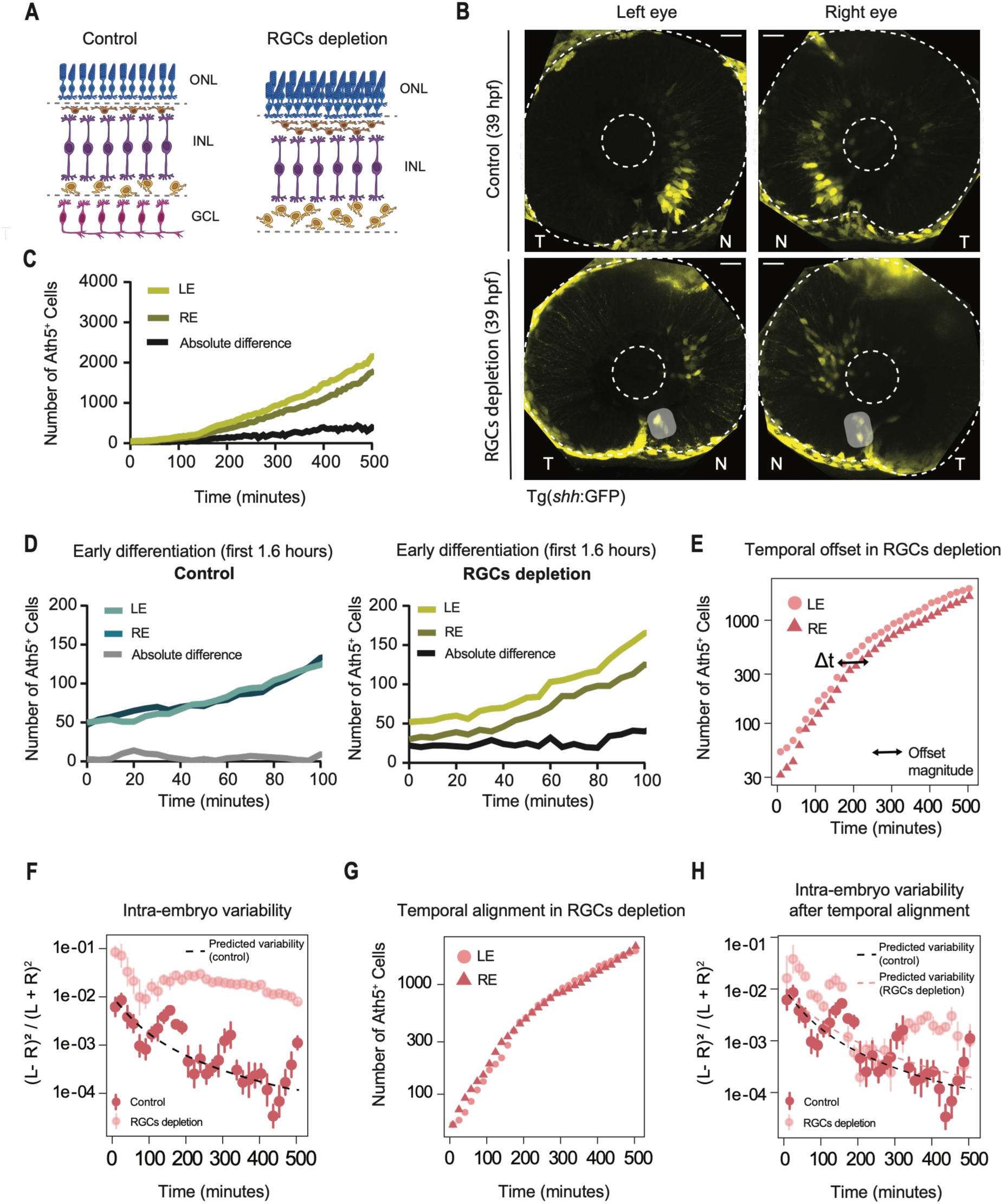
RGC depletion increases stochasticity of early Shh signalling and reduces bilateral matching of neurogenesis onset. (A) Schematics of control (left) and RGCs-depleted (right) retinas. Control retinas contain all five neuronal types distributed across the retinal ganglion cell layer (GCL), inner nuclear layer (INL) and outer nuclear layer (ONL). In RGCs-depleted retinas, the retinal ganglion cells (RGCs) are absent. (B) Representative Airyscan confocal images of Tg(*shh*:GFP) expression in paired left and right eyes at 39 hpf in a control embryo (top row) and RGCs-depleted embryo (bottom row). In controls, Shh expression is similar at the nasal pole. In RGCs-depleted embryos, Shh expression is sparse and more random between paired eyes. N, nasal pole; T, temporal pole. Dashed lines indicate the retinal outline and lens. Scale bars: 20 µm. (C) Quantification of Ath5^+^ cell emergence over time in the left (light green) and right (dark green) eyes of a representative RGCs-depleted embryo. The curves diverge progressively, with the absolute difference (black line) increasing over time. (D) Ath5^+^ cell emergence during early neurogenesis (first 100 minutes) in a representative control embryo (left) and a representative RGCs-depleted embryo (right). In controls, left-eye (light teal) and right-eye (dark teal) curves are closely matched from onset, with small absolute differences (grey line) between eyes. In RGCs-depleted embryos, left (light green) and right (dark green) eye curves diverge from the start, with larger absolute differences (black line) than in controls and increasing throughout the early period. (E) Ath5^+^ cell counts over time for the representative RGCs-depleted embryo shown in C, plotted on a semi-log scale. The horizontal offset (black arrow) between the left (circles) and right (triangles) eye curves indicates a delay in neurogenesis onset between the two eyes of the same embryo. (F) Relative intra-embryo variability (Left − Right)² / (Left + Right)^2^ over time for RGCs-depleted embryos (light red dots) and controls (dark red dots). The black dashed line indicates the model-predicted relative variability for the control condition, corresponding to the average variability over 100 samples from the posterior parameter distribution for each eye. RGCs-depleted embryos display substantially higher variability compared to both controls and the model prediction. (G) Temporal alignment of the representative RGCs-depleted embryo shown in C and E, after correcting for the estimated onset offset Δt. Following alignment, the left-eye (circles) and right-eye (triangles) curves overlap more closely, indicating that, after correction, differentiation dynamics remain largely conserved. (H) Relative intra-embryo variability (Left − Right)²/(Left + Right)^2^ over time after temporal alignment for RGCs-depleted embryos (light red dots) and controls (dark red dots). The black and red dashed lines correspond to the average relative variability over 100 samples from the posterior parameter distribution for each eye of the control condition and RGCs-depleted condition. Time is shown in minutes from the onset of differentiation (t = 0 defined as the time point at which the first eye of the pair reaches 50 Ath5⁺ cells). Scale bars: 20 µm.

To test whether indeed early Shh signalling was more variable between the two eyes of the same embryo after RGC depletion, we monitored its expression in fixed samples using the reporter transgenic line Tg(*shh*:GFP) in combination with the Tg(*ath5*:gap-RFP). In control embryos, Shh expression became detectable between 36 hpf and 37 hpf, initially emerging at the nasal pole (Fig. 6B). In contrast, in RGCs-depleted embryos, Shh onset was delayed being detected at 39 hpf and remained markedly reduced during early neurogenesis (Fig. 6B and Fig. S5A). Further, at 39 hpf, Shh expression between the two eyes was more spatially scattered and also more variable in RGC-depleted embryos than in controls (Fig. 6B), consistent with the reduction in early Shh-producing cells. Between 39 and 56 hpf, Shh expression in RGC-depleted retinas was sparser and more spatially heterogeneous than in controls but still followed the characteristic poles-to-mid-retina propagation pattern (Fig. S5A). At 56 hpf, Shh expression was restricted to the INL, as opposed to the RGC layer and INL in controls, consistent with ACs being the sole remaining intra-retinal source (Fig. S5A).

Having confirmed that intraretinal Shh onset was delayed and its expression reduced and more scattered in the Ath5 morphants, we monitored neurogenic wave onset and propagation by imaging the Tg(*ath5*:H2B-mNeonGreen) transgenic line and quantifying the number of emerging Ath5 positive cells over time (N = 3 embryos).

Indeed, we found that neurogenesis onset was more variable between the two eyes of the same embryo in Ath5 depleted embryos than in controls and in Ath5/Ptf1A depleted embryos (Fig. 6C,D and Fig. S5B). These differences emerged at neurogenesis onset and were not corrected during subsequent wave propagation but persisted over time. Consistent with this, the temporal offset between paired eyes (Δt; defined as the time shift required to align the Ath5 differentiation curves of the left and right eye; Fig. 6E) was increased in RGC-depleted embryos (mean absolute Δt = 53.89 min, N = 3 embryos, see SI) compared to controls (mean absolute Δt = 7.92 min, N = 4 embryos, see SI). Within this offset, both eyes increased Ath5 positive cell numbers at comparable rates (Fig. S5C), indicating conserved differentiation dynamics within embryos and leading to the amplification of initial differences over time (Fig. 6C,F and Fig. S5B,E). The spatial pattern of wave propagation followed the characteristic poles- to-mid-retina progression (Fig. S5D). Together, this indicated that once neurogenesis was underway, propagation proceeded with conserved dynamics in each retina independently of the initial offset.

In line with the experimental data, the model predicted that if each eye propagates independently from its own neurogenesis onset point, the spatiotemporal pattern of propagation should nevertheless be conserved, but with a temporal offset reflecting differences in neurogenesis onset timing between the two eyes. When the dynamics of each eye were rescaled to account for the offset of neurogenesis onset, the variability between the two eyes was greatly reduced, approaching the levels observed in the control condition (Fig. 6G, H). This strongly suggested that the increased between-eye variability in RGC-depleted embryos originated from stochastic differences in onset timing rather than propagation dynamics. Together, these results imply that once neurogenesis is underway, propagation proceeds independently and with conserved dynamics in each retina.

## Discussion

In this study, we investigated the onset and propagation of neurogenesis in the vertebrate retina, using the teleost zebrafish as a model. Its transparency and small size allowed quantitative 3D imaging over prolonged stages of neurogenesis. It further enabled the simultaneous imaging of both eyes within the same embryo, a prerequisite for the analysis of wave dynamics within embryos and comparison between embryos. The resulting quantitative framework enabled us to develop a stochastic model of wave propagation with the goal to distinguish between alternative scenarios for the regulation of neurogenesis onset and propagation between eyes.

We find that both the first Ath5 positive as well as the second Vsx1 positive neurogenic wave initiate at the nasal pole. After this nasal initiation, neurogenic cells soon emerge at temporal locations, and the wave fronts meet at mid-retinal positions. This contrasts with previous work that suggested a more direct nasal to temporal wave propagation (23, 24, 28, 33–35). These different interpretations most likely result from different methodologies used. It is possible that the previous 2D analyses of fixed samples were more prone to capture the dominant nasal wavefront, while the less prominent temporal one was harder to detect. Thus, this and other studies (57) show that using *in vivo* 3D imaging approaches at high time and spatial resolution over extended time periods can improve the understanding of spatiotemporal developmental patterning. We speculate that the short time interval between neurogenic onset at nasal and temporal poles is due to their close proximity, separated only by the choroid fissure, which would allow signals originating in the central brain (see below) to reach the two poles at similar timepoints.

When comparing wave dynamics across embryos, we find that these are generally conserved. However, the poor resolution of our initially used membrane and cytosolic markers restricted interpretation of data to qualitative observations. For a deeper understanding of neurogenesis wave initiation and propagation within and between embryos, we thus developed a new toolset combining a novel Ath5 nuclear reporter line, Tg(*ath5*:H2B-mNeonGreen) with a 3D IMARIS-based nuclei detection pipeline. This allowed us to follow onset and propagation of the first Ath5 positive neurogenic wave at single-cell resolution over time and in space. With this pipeline we were able to quantitatively characterise the differentiation dynamics and make systematic comparisons between embryos. We find that the main sources of inter-embryo variability correspond to the onset timing and the spatial propagation of the differentiation wave. These differences are most likely due to variations in developmental timing between individuals.

When comparing emerging cell numbers and spatial progression of neurogenic cells between the two eyes of the same embryo, however, we find that here neurogenesis onset and propagation are considerably more closely matched than between embryos. At first sight, this finding could suggest that embryos actively control the progress of neurogenesis possibly via eye-to-eye communication. We find, however, that despite their similarity, the two eyes of the same embryo show small differences in cell numbers over time. To be able to understand whether these differences can arise from the intrinsic stochasticity of the differentiation process, we developed a model in which differentiation decisions are stochastic and cell-autonomous, with the probability of differentiation depending on the position along the naso-temporal axis. This stochastic nature of individual differentiation decisions in our model is consistent with earlier work showing that retinal progenitors produce clones of variable size and composition (40, 58). Together, these studies suggest that individual fate decisions carry an inherent stochastic component.

After fitting our model to the experimentally observed cell counts, it suggested that the observed parallel progression of Ath5 dependent neurogenesis could be explained by a shared onset of neurogenesis alone, without a need for an ongoing communication between the eyes. The remaining differences between eyes are consistent with the idea that each retina propagates independently through neurogenesis and reaches the degree of bilateral matching observed.

Based on previous data (24, 31, 33–35, 59), we speculated that intraretinal Shh produced by retinal ganglion cells (RGCs) (36) and amacrine cells (ACs) (34) could play a major role in wave propagation. When we eliminated the majority of these cells that serve as intra-retinal Shh sources by suppressing their emergence (26, 54–56), neurogenesis onset was delayed but started at the same time in both eyes and from then remained closely matched over time albeit with slightly higher variability than in controls. What changed in this condition was that wave progression did not follow the typical pole-to-mid-retina pattern, but neurogenic cells emerged across the retina from early neurogenesis stages. The modest increase in cell number variability between the two eyes of an embryo could be accounted for by the model assuming a flatter differentiation profile while keeping the average differentiation time measured across control embryos. Thus, the bilateral similarity observed for the first neurogenic wave is generally robust even when intra-retinal Shh is removed and spatial organisation of neurogenesis is altered. This argues that spatial patterning of the wave and the bilateral symmetry that is observed are separable traits. Currently, we do not know what signals or other factors ensure that despite patterning disruption neurogenesis still shows bilateral similarity.

To directly test whether indeed a shared onset of neurogenesis is a prerequisite for bilateral matching during wave propagation, we depleted Ath5, which suppresses the emergence of RGCs (26, 55). As noted above, RGCs and ACs are the main producers of early intra-retinal Shh (32, 34), with RGCs contributing in significantly bigger proportion at early neurogenesis stages (32, 33). When RGCs do not emerge, neurogenic division behaviour at these early neurogenic stages shifts from the production of one Shh-generating RGC and one PR to mainly symmetric PR-PR divisions with only few early divisions generating PR-AC offspring (26). As in this scenario ACs are the only remaining cells that produce intra-retinal Shh, this strong reduction, specifically during the initiation of neurogenesis, was postulated to make early wave propagation more susceptible to stochastic fluctuations between eyes. At later stages, once sufficient ACs are produced, propagation was expected to normalise.

Indeed, using this regime neurogenesis onset became more variable between the two eyes of the same embryo when compared to controls. These differences in neurogenic outcome persisted over time rather than being corrected during neurogenesis propagation. Within this offset, however, both eyes propagated with conserved dynamics and spatial patterning from nasal and temporal to mid-retina positions was conserved. When each eye’s dynamics were rescaled and aligned to a shared onset, bilateral matching was restored almost to levels seen in controls. This experimental outcome matched the model’s prediction, meaning that indeed the simultaneous timing of neurogenic onset could be a major determinant of bilateral similarity.

While we are aware that morpholino use has been debated in the field and off-target effects and compensation mechanisms were in some instances reported (60–62), we are confident that the reagents used here are well-validated (17, 26, 54–56). They further have been shown to reproduce the corresponding mutant phenotypes across multiple independent studies (17, 26, 55, 56, 61, 63).

In sum, the outcome of our combination of model and experiment suggests that no inter-retinal communication is necessary for maintaining bilateral similarity between eyes. Despite these findings, we do not exclude that such communication exists. This additional communication may play a role when perturbations need to be buffered or for fine-tuning of bilateral matching under challenging developmental conditions. It was for example shown that when one eye is physically reduced in size, the other eye can show delayed neurogenesis onset (64). Even though this previous study did not directly compare neurogenesis in the two eyes to each other in a quantitative manner, this finding could mean that some form of inter-organ communication exists that allows the two eyes to sense and respond to each other’s developmental state (64). Similar buffering mechanisms have also been proposed in other paired organ systems, such as the zebrafish otic vesicles, where bilateral similarity can recover after asymmetric perturbations affecting tissue growth and size in a single ear. Interestingly, here recovery appears to rely on organ autonomous compensatory mechanisms rather than interactions between the two otic vesicles (14).

To resolve the question of whether inter-eye communication exists after neurogenesis has started, future studies will have to directly perturb one eye while monitoring the other, for example by repeating the cell depletion experiments mentioned above or by inducing cell death in a spatiotemporally controlled manner. It would further be insightful to move beyond cell number quantification as presented in this work, towards an approach that allows to analyse cells entering neurogenesis in direct spatiotemporal relation to each other. This would determine whether neighbouring cells influence each other’s decision to differentiate. Deep learning-based cell tracking approaches, which are rapidly becoming accessible and more reliable, will likely be key to understand which cells enter neurogenesis, when, and where at the necessary scale and resolution.

Another question emerging from this study is what factors initially trigger the shared neurogenesis onset. Central brain-derived Shh upstream of Fgf signalling is an attractive and likely candidate route as it has already been shown that these factors are necessary (24) and, importantly, also sufficient (35, 45, 59) for neurogenesis to start. It is tempting to speculate that these factors could derive from the central brain and reach both eyes simultaneously via diffusion or via cell-to-cell spreading, assuming that eyes are symmetrically positioned at equal distance from the source. Interference with these candidates in a spatiotemporally defined manner will be key to dissect their exact contributions. Additional signals originating from the brain or surrounding tissue most likely also play a role in neurogenesis onset and propagation. Single cell as well as spatial transcriptomics (to compare the two eyes at the same stage) will be essential to uncover these additional signals that could fine-tune symmetric neurogenesis across waves.

The findings presented here were only possible due to the combination of quantitative live imaging and theoretical modelling as it allowed us to move from observation to testable experimental frameworks. However, the model has limitations as we only follow neurogenesis for the initial 500 minutes. We assume that with continued development all cells will differentiate, which in turn would lead to a natural decrease in variability, which we do not account for. It will be interesting to probe whether this final progenitor depletion can by itself lead to neuronal patterning correction in perturbed conditions or whether additional error correction mechanisms would be needed.

It will further be interesting to test whether the model can be generalised to eyes of different sizes, given that the intra-embryo variability depends on the total cell number. Testing this in differently sized organisms would reveal whether there are limits to achieving symmetry through the stochasticity of dynamics alone in the retina and beyond.

## Materials and methods

### Zebrafish husbandry

Wildtype zebrafish (*Danio rerio*) strains (AB and TL) and transgenic lines were maintained under standard conditions at 26 °C. Embryos were staged in hours post-fertilisation (hpf) according to (65). Embryos of undetermined sex were used between 24 and 52 hpf. For experiments, embryos were raised at 28 °C in E3 medium supplemented with 0.2 mM 1-phenyl-2-thiourea (PTU; Acros Organics, 10107703) from 8 hpf to prevent pigmentation. Methylene blue was added to inhibit fungal growth. Medium was replaced daily. For live imaging, embryos were anaesthetised in 0.04% tricaine methanesulfonate (MS-222; Pharmaq, 1004671) in E3 and maintained under anaesthesia throughout acquisition.

All procedures were conducted in accordance with institutional and national guidelines under DGAV (Portugal) licensing, in compliance with EU Directive 2010/63/EU and Portuguese Decree Law 113/2013.

### Transgenic lines

The Tg(*ath5*:gap-RFP) line was used to label the membranes of Ath5^+^ progenitors and Ath5^+^ neurons (48). Tg(*ath5*:H2B-mNeonGreen) (this study, see below) was used to label the nuclei of Ath5^+^ progenitors and neurons. Tg(*vsx1*:GFP) was used to label the cytoplasm of Vsx1^+^ progenitors and neurons (66). Tg(*lama1:lama1*-mKate2) was used to visualise the retinal basal lamina and retinal outline (53).

### Plasmid construction and line generation

#### Tg(*ath5*:H2B-mNeonGreen)

This construct contains an H2B-mNeonGreen fusion protein under the control of the *ath5* regulatory sequence and was assembled using a Gateway/Tol2 cloning strategy. Prior to stable line generation, construct functionality was validated in two steps. First, the plasmid was injected into Wildtype AB embryos to confirm nuclear fluorescence expression in Ath5-expressing territories. Second, the plasmid was injected into Tg(*ath5*:gap-RFP) embryos, and the expression pattern was assessed by fluorescence microscopy, confirming co-expression with the existing Ath5 reporter (Fig. 3A). Fluorescence was detected in known Ath5-expressing tissues, including the retina and olfactory epithelium from 24hpf onwards.

For stable transgenic line generation, the plasmid DNA (25 ng/µL) was co-injected with Tol2 transposase RNA (50 ng/µL) and PCS2+ mKate2-ras RNA (25 ng/µL) (44) into one-cell-stage embryos. Injected embryos were screened for mosaic fluorescence expression. Fluorescence-positive F0 founders were raised to adulthood and outcrossed to Wildtype fish. F1 progenies were screened for stable germline transmission by fluorescence microscopy and positive individuals were incrossed to establish the stable Tg(*ath5*:H2B-mNeonGreen) line.

### Morpholino experiments

All morpholinos were purchased from GeneTools.

The following amounts of morpholino were injected into the yolk at the one-cell stage:

ng of Ath5-MO 5’-TTCATGGCTCTTCAAAAAAGTCTCC-3’ (55);
ng Ptf1a-MO1 5’-CCAACACAGTGTCCATTTTTTGTGC-3’ (54);
ng Ptf1a-MO2 5’-TTGCCCAGTAACAACAATCGCCTAC-3’ (54).
ng p53 MO 5’-GCGCCATTGCTTTGCAAGAATTG-3’ (67) were added to prevent increased apoptosis.

To exclude off target effects, for all experiments, control embryos were injected with a scrambled morpholino

5’-CCTCTTACCTCAGTTACAATTTATA-3’ at 0.5 ng per embryo, together with the p53 morpholino. These embryos showed control morphology.

Morpholino knockdown efficiency was assessed after image acquisition. Ath5-MO and Ath5-MO; Ptf1a-MO embryos were screened for absence of the optic nerve, and embryos displaying an optic nerve were excluded from subsequent analysis. To validate Ptf1a knockdown efficiency, Ptf1a-MO-injected embryos were fixed after live imaging and immunostained for HuC/HuD. Embryos displaying HuC/HuD-positive interneurons within the inner nuclear layer were excluded from the analysis.

### Whole-mount immunofluorescence

Zebrafish embryos were dechorionated and fixed overnight at 4°C in 4% paraformaldehyde (Thermo Fisher Scientific, 043368-9 M) in PBS. Samples were washed three times in PBST (PBS containing 0.2% Triton X-100; VWR, 28817295) for 10 minutes each at room temperature on a shaker, then rinsed in H2O to remove residual salts. For permeabilisation, embryos were incubated at -20°C in cold acetone for 15 minutes. After rehydration in PBS with 0.5% Triton, embryos were incubated in a blocking solution containing 10% Donkey serum in PBS with 0.2% Tween for 2 hours at room temperature.

Embryos were incubated with primary antibodies diluted in antibody solution for 64 hours at 4°C. The primary antibodies used were anti-HuC/HuD (1:250; Invitrogen, A-21271) and anti-Zn5 (1:20; Zirc, ZDB-ATB-081002-19). Embryos were washed five times for 20 minutes each with PBS containing 0.5% Triton. They were then incubated with the secondary antibodies in the antibody solution (antibodies diluted 1:200) and DAPI at 1:1000 (Thermo Fisher Scientific) for 2 days at 4°C.

The following secondary antibodies (Thermo Fisher Scientific) were used at 1:200: Alexa Fluor 647 goat anti-mouse (A-21236) and Alexa Fluor 488 donkey anti-rabbit (A-21206).

RFP-Booster (rba594, Chromotek) and GFP-Booster (gba488, Chromotek) were added at this step at 1:200 to enhance RFP and GFP fluorescence, respectively.

Embryos were washed four times for 15 minutes each in PBS with 0.5% Triton and stored in PBS with 0.02% sodium azide at 4°C until imaging.

### Confocal microscopy

Fixed, whole-mount immunostained embryos were imaged using a Zeiss LSM980 Airyscan2 inverted point-scanning confocal microscope equipped with two photomultiplier tube (PMT) detectors and one gallium arsenide phosphide (GaAsP) detector, using the 40x/1.1 C-Apochromat water immersion objective (Zeiss). Embryos were mounted in 0.7% low-melting agarose in 35-mm glass-bottom dishes (MatTek Corporation) and imaged at room temperature. Z-stacks spanning the entire retina were acquired with a z-step size of 1 µm. Image acquisition was performed using ZEN Blue v3.3 software (Zeiss).

### *In vivo* light-sheet fluorescence microscopy (LSFM)

Embryos were staged and manually dechorionated at 24hpf. Before mounting, embryos were screened for fluorescence. After screening, dechorionated embryos were mounted in 1-mm glass capillaries in 0.6% low-melting agarose as previously described in (68). The light-sheet chamber was filled with media, composed of E3-water medium containing 0.2 mM PTU (Sigma) and 0.01% MS-222 (Sigma). Imaging started at 28 hpf and was performed for 24 hours, until 52 hpf, encompassing the onset and propagation of both the Ath5 and Vsx1 neurogenic waves. Imaging was performed using a Zeiss Lightsheet Z.1 microscope equipped with two PCO Edge 4.2 sCMOS cameras (max 30 fps with 2048 x 2048 pixels - pixel size 6.5 µm), with a 20X/1.2 Zeiss Plan-Apochromat water immersion objective. Imaging was performed at 28°C. Z-stacks spanning the entire retinal neuroepithelium (160 µm) were acquired with 1-µm optical sectioning every 5 minutes for 24 hours in double-sided illumination mode. Up to two embryos were imaged per imaging session. The system was operated using the ZEN 3.1 software (black edition).

### Quantitative analysis of Ath5 and Vsx1 neurogenic onset and propagation

#### Movie preprocessing

Light-sheet imaging datasets of Ath5 and Vsx1 neurogenic dynamics were cropped to include only the retinal region of interest. Signals from the two illumination sites were fused in ZEN Black (Zeiss) to maximise signal recovery across the retinal tissue. Images were subsequently downscaled two-fold in the X and Y dimensions and processed in Fiji for 2D analyses or IMARIS for 3D analyses.

#### Retinal straightening for 2D kymograph generation

To analyse the spatiotemporal dynamics of Ath5 and Vsx1 expression (Fig. 2A-A’’ and S1), light-sheet movies were projected along the z-axis using maximum-intensity projections of fluorescence intensity at each time point. To compensate for tissue growth and sample drift during the 24 h imaging period, movies were registered using the Fiji StackReg plugin (Affine transformation; https://imagej.net/plugins/stackreg). The final time point, corresponding to the maximum retinal thickness reached during the imaging period, was used as the reference frame for registration. A segmented line was manually drawn along the naso-temporal arc of the retina, spanning from the nasal to the temporal pole. The line width was adjusted to encompass the full retinal thickness, and the same segmented line was applied to all time points and channels (Ath5 and Vsx1) within each dataset. The retinal region defined by the segmented line was subsequently straightened using the Reslice plugin in Fiji (https://imagej.net/imaging/z-functions#stack-reslice), thereby generating 2D kymographs representing spatiotemporal fluorescence dynamics.

#### Quantification of Ath5 and Vsx1 neurogenic onset from 2D kymographs

To quantify Ath5 and Vsx1 neurogenic onset (Fig. 2B, C), fluorescence intensity profiles were extracted from the retinal kymographs generated, as described above. Kymographs were rotated by 90° so that developmental time was represented along the y-axis. The delay between Ath5 and Vsx1 onset was calculated as the difference between the onset times of the two reporters within the same retina.

#### Quantification of Ath5 positive cells in fixed Airyscan z-stacks

Embryos were fixed at 30 hpf (Fig. S2), and retinal z-stacks were acquired by Airyscan confocal microscopy (see Imaging). Individual Ath5^+^ cells were identified based on Tg(*ath5*:gap-RFP) expression and counted throughout the entire retinal volume using the Cell Counter plugin in Fiji (https://imagej.net/plugins/cell-counter). Z-stacks spanned the entire retina (> 120 µm per retina; optical sections acquired every 1 µm). Cells appearing in consecutive optical sections were tracked through the z-stack and counted only once. Left and right eyes were quantified independently.

#### IMARIS-based 3D detection of Ath5 positive nuclei

Retinal neurogenesis dynamics were analysed *in vivo* using light-sheet imaging of Tg(*ath5*:H2B-mNeonGreen) and Tg(*lama1:lama1*-mKate2) double transgenic embryos. Prior to spot detection, datasets were manually corrected for drift in IMARIS by tracking a reference position within the ciliary marginal zone (CMZ) throughout the movie and applying translational corrections in the x, y and z dimensions to all objects. Ath5 positive nuclei were subsequently detected in 3D using the Spot detection module in IMARIS (v10.2; Bitplane, Oxford Instruments) applied to the Tg(*ath5*:H2B-mNeonGreen) channel. The Tg(*lama1:lama1*-mKate2) reporter was used to define the retinal outline and exclude nuclei outside the retinal tissue. Spot detection was performed using a spot diameter of 3 µm. Fluorescence intensity thresholds were determined on a representative subset of time points and applied consistently throughout the entire time series. Detected spots were manually inspected and curated blind to experimental conditions. Manual corrections were restricted to clear detection artefacts, including missed, false-positive, or merged spots.

#### Quantification of Ath5 positive cell emergence dynamics

Time-resolved spot coordinates and counts were exported from IMARIS for downstream analysis in GraphPad (v9.4.0 (673)) and R 4.5.3. For each retina, the total number of Ath5 positive nuclei within the retinal outline was quantified at each time point during the first 500 min of imaging. For comparisons within and between embryos, curves were normalised to the onset of neurogenesis, defined as t = 0 at the time point when the first eye of the pair reached 50 detected Ath5 positive cells. Intra-embryo absolute differences in Ath5 positive nuclei number were calculated at each time point by subtracting the count of the right eye from that of the left eye of the same embryo.

#### Quantification of Ath5 positive cell emergence rates

To measure Ath5 positive accumulation changes over time, the rate of Ath5 positive cell emergence was calculated as the change of the cumulative Ath5 positive cell count with respect to time between consecutive imaging frames:

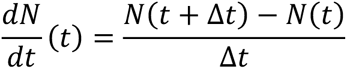

where *N(t)* represents the number of detected Ath5 positive nuclei at time *t* and Δ*t* corresponds to the imaging interval (5 min). Differentiation rate profiles were calculated independently for each retina.

#### Data, Materials, and Software Availability

The R scripts used for Bayesian inference and stochastic simulations have been deposited in the GitHub repository (https://github.com/al-sina/bilaterality). All study data are included in the article and/or SI Appendix.

#### Declaration of generative AI and AI-assisted technologies in the writing process

During the preparation of this work, the authors used GPT-5.5 and Claude 4.6 to edit the manuscript for spelling, grammar, and clarity. The authors critically reviewed and edited the output of these tools and assume full responsibility for the content of this publication.

## Acknowledgments

We thank all members of the Cell Biology of Tissue Morphogenesis laboratory for lively project discussions. We thank Clara Mestre, Diana Carrasqueira, and Renata Cunha for experimental support, Jaroslav Icha for cloning the *Ath5:H2B-mNeonGreen* plasmid and Robert Haase for helpful discussions on image analysis and for recommending the Fiji plugins used for retinal straightening. We thank Elisa Nerli for valuable discussions during the conception and early development of the project, and José Freitas for valuable project discussions. We thank Pablo Sartori and Tiago Paixão for fruitful input on quantitative analysis. We thank Lucrezia Ferme, Jake Cornwall Scoones and Michel Cayouette for constructive comments on the manuscript. We are grateful to the Advanced Imaging and Aquatic Facilities at the Gulbenkian Institute for Molecular Medicine (GIMM, formerly IGC) and to Davide Accardi at the Advanced Bioimaging and Biooptics Facility at Champalimaud Research for technical support.

## Author Contributions

D.G.-M.: Conceptualisation, data acquisition and curation, investigation, methodology, validation, visualisation, formal analysis, writing - original draft. A.A.: Conceptualisation (theoretical model), software, data curation, methodology (theoretical model), validation, formal analysis, writing - supplementary theory, writing - original draft. G.S.: Conceptualisation, supervision (theoretical model), writing - original draft. C.N.: Conceptualisation, supervision, project administration, funding acquisition, writing - original draft.

## Extended data figures and figure legends

**Figure S1.**
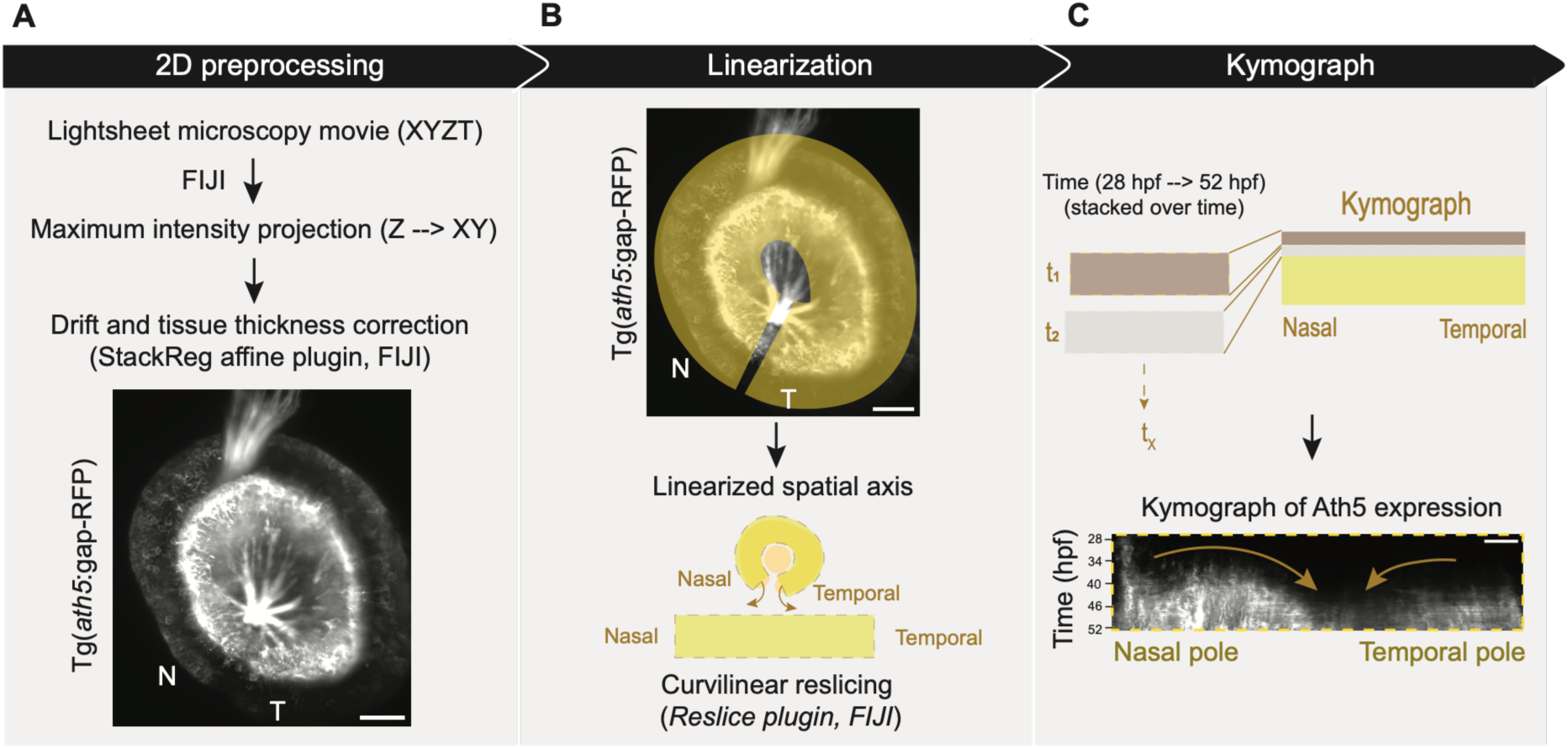
Image processing pipeline for 2D kymographs. (A) Preprocessing of light-sheet microscopy movies (XYZT). For each time point, a maximum intensity projection along the z-axis was computed in FIJI. The resulting movie was then corrected for drift and tissue thickening over time using the StackReg plugin (affine transformation mode) in FIJI. (B) Spatial linearization of the curved retinal tissue. A curved, segmented line (yellow) was applied along the nasotemporal arc of the retina, from the nasal pole (N) to the temporal pole (T), avoiding overlap between the two poles. The same segmented line was applied to both the Tg(*ath5*:gap-RFP) and Tg(*vsx1*:GFP) channels. The curved tissue was linearised using the Reslice plugin in FIJI, generating a flat spatial representation of the naso-temporal axis for each time point. (C) The Reslice plugin automatically generated a 2D kymograph by stacking the linearised spatial profiles from each time point (t_1_, t_2_, …t_x_) along the time axis. In the resulting kymograph, the x-axis represents position along the naso-temporal axis and the y-axis represents developmental time (28-52 hpf). A representative Ath5 kymograph is shown in greyscale. Scale bars: 20 µm.

**Figure S2.**
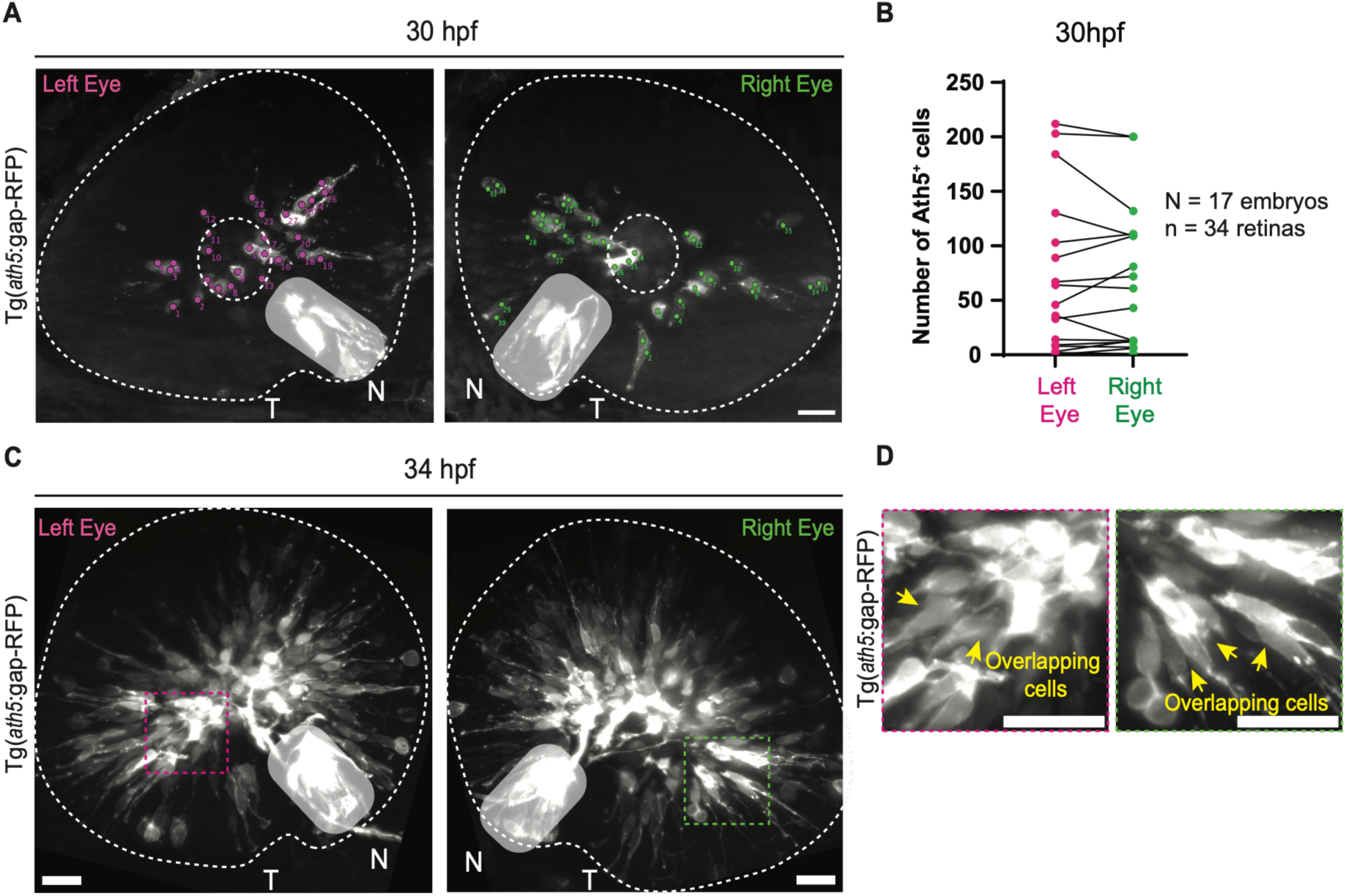
Manual quantification of Ath5^+^ cells during early retinal neurogenesis. (A) Representative maximum intensity projections of left and right retinas from a single embryo at 30 hpf, showing Tg(*ath5*:gap-RFP) expression. Dots indicate Ath5^+^ cells (magenta, left eye; green, right eye) annotated using the Cell Counter plugin in FIJI (see Methods). Dashed lines represent the retinal outline and the position of the lens. (B) Paired quantification of Ath5^+^ cell number at 30 hpf across embryos. Each pair corresponds to the left (magenta) and right (green) eyes of a single embryo. Black lines connect the two eyes of the same embryo (N = 17 embryos, n = 34 retinas). (C) Representative maximum intensity projections of left and right retinas of the same embryo at 34 hpf. Compared to earlier stages, Ath5^+^ cells are densely packed and overlapping. Dashed white lines represent the retinal outline and the position of the lens. Dashed regions of interest (ROIs) (magenta, left eye; green, right eye) indicate the regions shown at higher magnification in (D). (D) Close-up views of the ROIs in (C). Yellow arrows indicate regions of densely overlapping Ath5^+^ cells with ambiguous boundaries, illustrating the limitations of manual quantification at later stages. N, nasal pole; T, temporal pole. The regions highlighted in grey correspond to the ciliary margin zone (CMZ). Scale bars: 20 µm.

**Figure S3.**
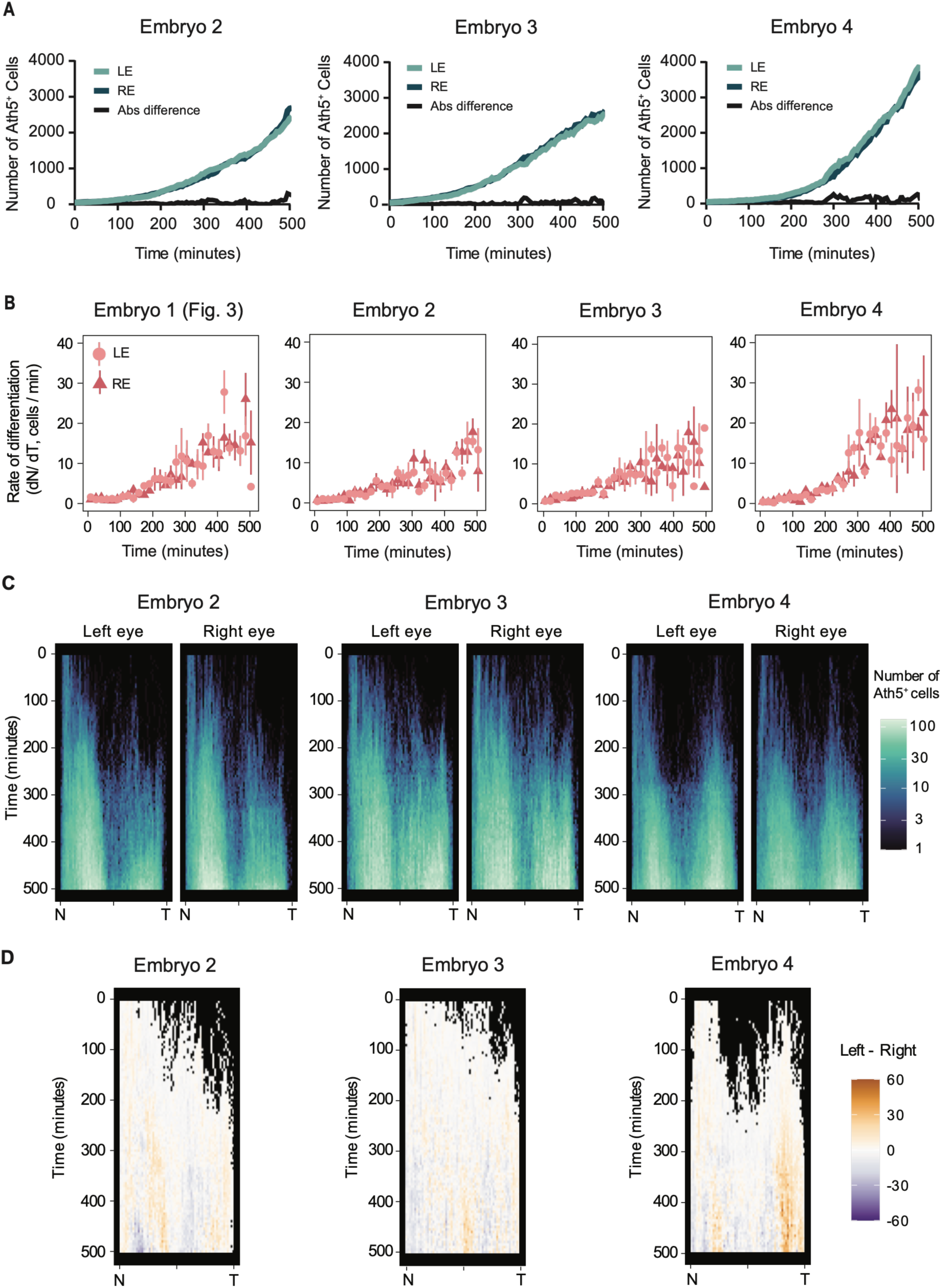
Ath5^+^ cell number emergence, differentiation rates and spatiotemporal propagation in control embryos. (A) Ath5^+^ cell number over time in the left (LE, light teal) and right (RE, dark teal) eyes of control embryos, with the absolute difference between eyes shown in black. (B) Differentiation rates (dN/dT, cells/min) over time for the left (LE, red circles) and right (RE, dark red triangles) eyes of control embryos (N = 4 embryos). Embryo 1 corresponds to the representative embryo shown in Figure 3. (C) Spatiotemporal kymographs showing the number of Ath5^+^ cells per naso-temporal position over time for the left and right eyes in control embryos. The number of Ath5^+^ cells is colour-coded ranging from 0 (black) to 100 (white). (D) Intra-embryo difference kymographs (LE – RE) across the naso-temporal axis over time for embryos shown in (C). Differences are colour-coded from - 60 (purple, more cells in the right eye) to 60 (orange, more cells in the left eye), with white indicating no difference. Black regions reflect positions and time points at which Ath5^+^ cells are absent in both eyes. Time is indicated in minutes from the onset of differentiation (*t* = 0 defined as the time point at which the first eye of the pair reaches 50 Ath5^+^ cells). N, nasal pole; T, temporal pole.

**Figure S4.**
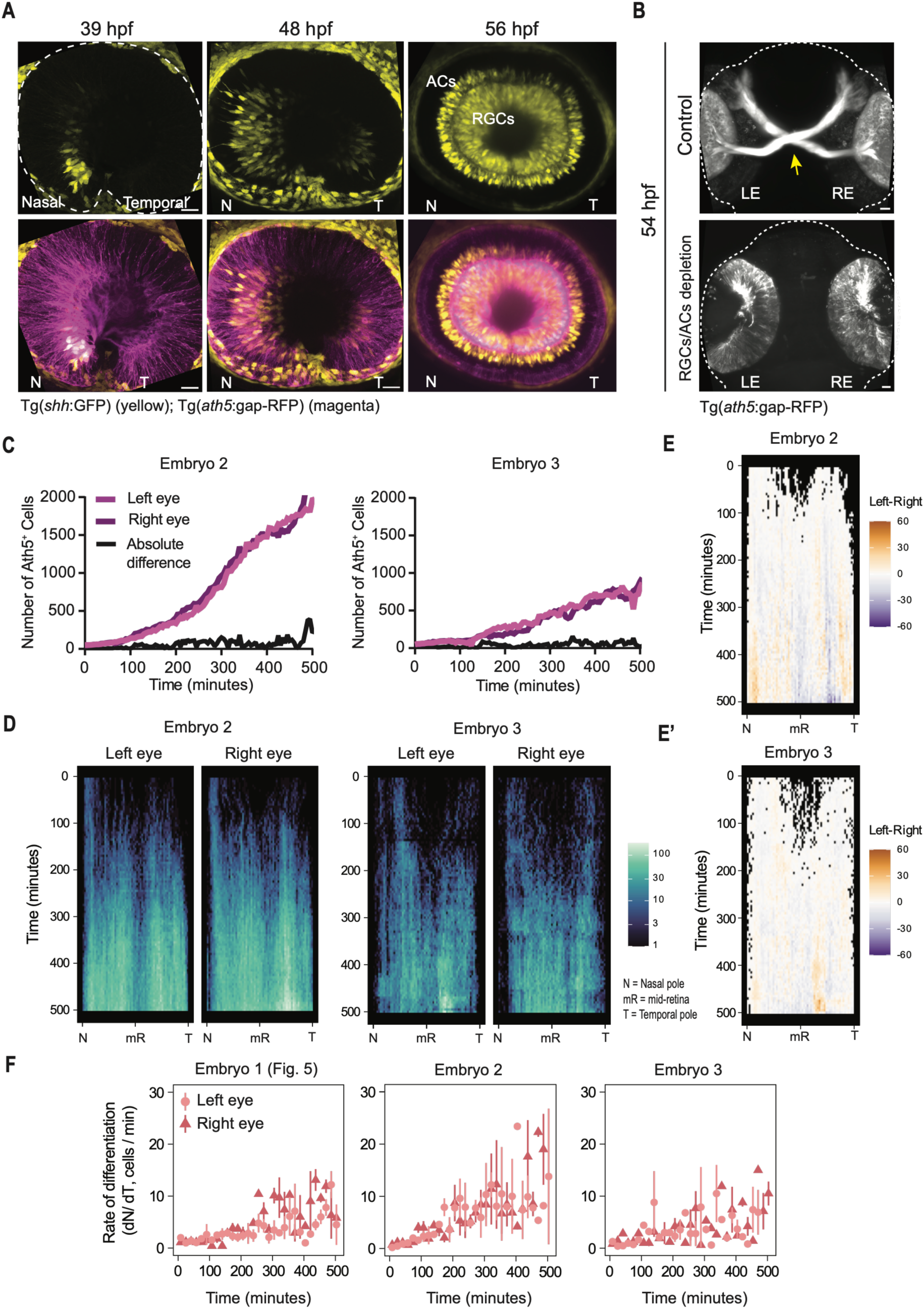
Reduced Shh signalling flattens Ath5^+^ wave propagation while preserving bilateral synchrony after RGCs/ACs depletion. (A) Representative maximum intensity projections of a control embryo at 39 hpf, 48 hpf and 56 hpf showing Tg(*shh*:GFP) alone (top row, yellow) and merged with Tg(*ath5*:gap-RFP) (bottom row; Ath5:gap-RFP, magenta; Shh:GFP, yellow). (B) Representative Tg(*ath5*:gap-RFP) images at 54 hpf in control (top) and RGCs/ACs depleted (bottom) embryos. In controls, RGC axons build the optic nerve and project through the optic chiasm (yellow arrow). In depleted embryos, the optic nerve is absent, confirming RGC depletion. LE, left eye; RE, right eye. Dashed lines indicate the head outline. (C) Quantification of Ath5^+^ cell emergence over time in left (light magenta) and right (dark magenta) eyes of RGCs/ACs-depleted embryos (Embryos 2 and 3). The black line represents the absolute intra-embryo difference between eyes. (D) Kymographs showing the Ath5^+^ cell number per naso-temporal position over time for the left and right eyes of the embryos shown in (C). The number of Ath5^+^ cells is colour-coded ranging from 0 (black) to 100 (white). (E-E’) Intra-embryo difference kymographs of Ath5^+^ cell numbers for Embryo 2 (E) and Embryo 3 (E’). Differences are colour-coded from -60 (purple, more cells in the right eye) to 60 (orange, more cells in the left eye), with white indicating no difference. Black regions indicate time points and positions at which Ath5^+^ cells are absent in both eyes. (F) Rate of Ath5^+^ cell differentiation (dN/dT, cells/min) over time for left (red circles) and right (dark red triangles) eyes across all three RGCs/ACs-depleted embryos. Embryo 1 corresponds to the representative embryo shown in Figure 5. Time is indicated in minutes from the onset of differentiation (*t* = 0 defined as the time point at which the first eye of the pair reaches 50 Ath5^+^ cells). N, nasal pole; mR, mid-retina; T, temporal pole. Scale bars: 20 µm

**Figure S5.**
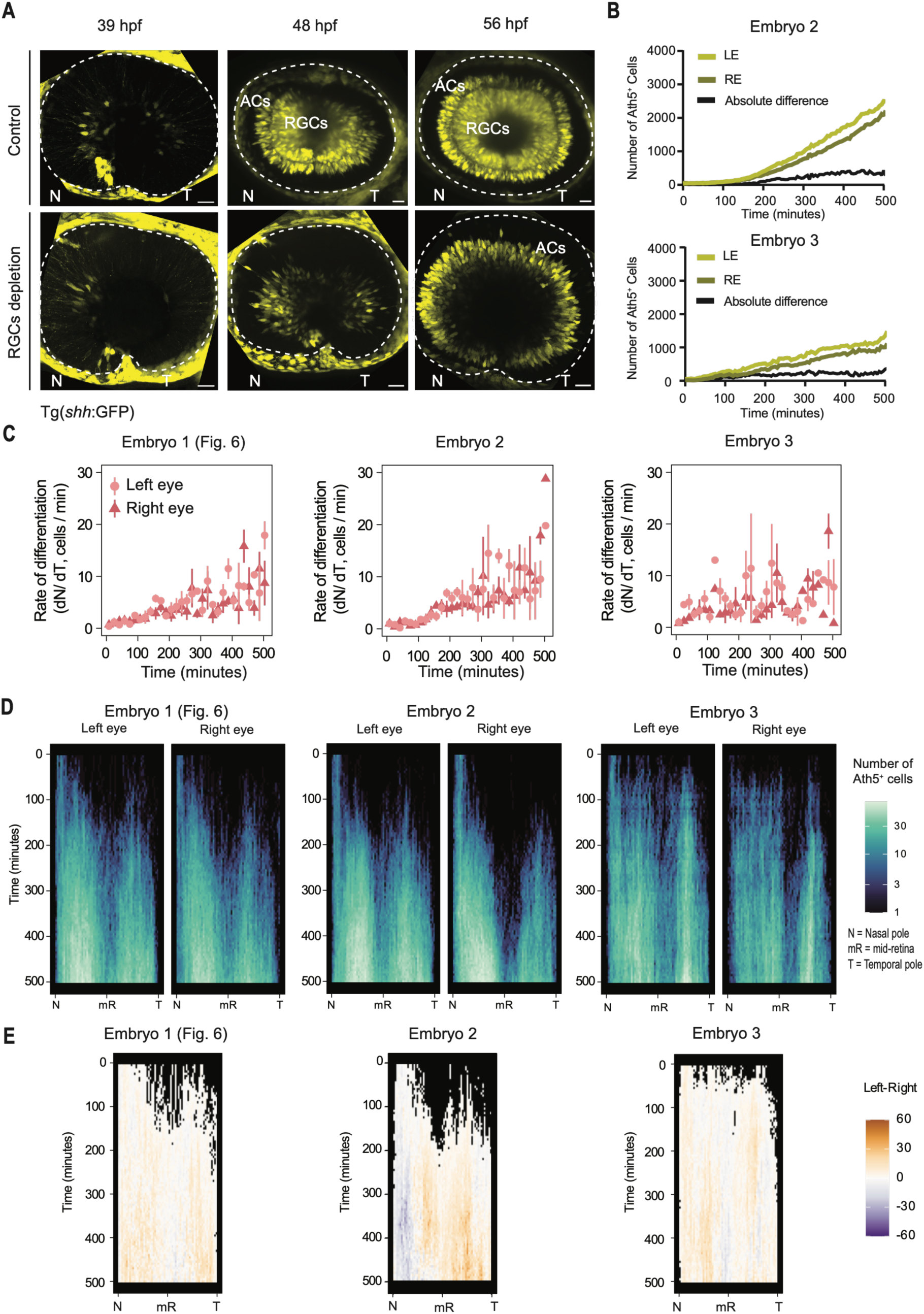
Reduced and asymmetric Shh signalling is associated with uncoupled bilateral neurogenesis after RGC depletion. (A) Representative images of Tg(*shh*:GFP) expression at 39 hpf, 48 hpf and 56 hpf in a control embryo (top row) and an RGCs-depleted (bottom row) embryo. In controls, Shh expression initiates at the nasal pole and at 39 hpf at the temporal pole and progressively propagates toward the mid-retina thereafter. From 48 hpf, two distinct layers are visible: the inner layer corresponding to retinal ganglion cells (RGCs) and the outer layer corresponding to the amacrine cells (ACs). In RGCs-depleted embryos, Shh onset is detected at the nasal pole at 39 hpf and at the temporal pole by 48 hpf. By 56 hpf, only a single layer is present in the INL, corresponding to ACs. N, nasal pole; T, temporal pole. Scale bars: 20 µm. (B) Quantification of Ath5^+^ cell emergence over time in left (light green) and right (dark green) eyes of embryos 2 (top) and 3 (bottom) from the RGCs-depleted condition, with absolute intra-embryo difference shown in black. Absolute differences are shown as black line. (C) Rate of Ath5^+^ cell differentiation (dN/dT, cells/min) over time for left (red circles) and right (dark red triangles) eyes of the three RGCs-depleted embryos. Embryo 1 corresponds to the embryo shown in Figure 6. (D) Quantitative spatiotemporal kymographs showing the number of Ath5^+^ cells per naso-temporal position over time for the left and right eyes of the three RGCs-depleted embryos. Embryo 1 corresponds to the representative embryo shown in Figure 6. The number of Ath5^+^ cells is colour-coded ranging from 0 (black) to 100 (white). (E) Intra-embryo difference kymographs (left - right) along the naso-temporal axis over time for embryos 1, 2 and 3. Differences are colour-coded from -60 (purple, more cells in the right eye) to 60 (orange, more cells in the left eye), with white indicating no difference. Black regions correspond to positions and time points at which Ath5^+^ cells are absent in both eyes. Time is indicated in minutes from the onset of differentiation (t = 0 defined as the time point at which the first eye of the pair reaches 50 Ath5^+^ cells).

## Supplementary movie legends

**Movie S1. Simultaneous live imaging of the Ath5 and Vsx1 neurogenic waves (related to Fig. 1C, D).**

Lightsheet movie of onset and propagation of the first and second neurogenic waves. The first neurogenic wave is visualized using Tg(*ath5*:gap-RFP) (cyan in the merged view), and the second neurogenic wave is visualized using Tg(*vsx1*:GFP) (magenta in the merged view). Embryos were imaged starting at 28 hpf in 5-minute intervals. Scale bar, 20 µm.

**Movie S2. Simultaneous live imaging of Ath5^+^ cells emerging within the retinal tissue (related to Fig. 3 B, C).**

**Part 1:** 3D view of the emergence of Ath5+ cells within the retinal tissue using Tg(*ath5*:H2B-mNeonGreen) (cyan). Retinal outline is labelled by Tg(*lama1:lama1*-mKate) (grey).

**Part 2:** Three-dimensional IMARIS spot detection of Ath5+ nuclei (FIRE LUT according to proximal-distal position) overlaid with Tg(*ath5*:H2B-mNeonGreen) signal.

**Part 3:** Three-dimensional IMARIS spot detection overlaid with Tg(*lama1:lama1*-mKate), which labels the retinal outline.

Embryos were imaged starting at 28 hpf with 5-minute intervals. Scale bar, 15 µm.

## Supplementary Theory

In this document we provide additional details and mathematical derivations of the analytical results presented in the main text.

### I. DATA PREPROCESSING

The IMARIS spot detection pipeline provides the spatial positions (*x, y, z*) of all nuclei detected in the retina at each time point *t*. To standardise the data collected across different eyes for subsequent analysis, we first reflect the *x* coordinate of right eyes so that both eyes share the same orientation. We then normalise the *x* and *y* coordinates by centering the data around the late-stage center of mass as

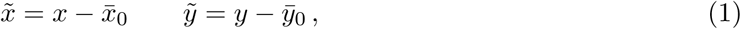

where (*x̅*_0_*, y̅*_0_) are the coordinates of the center of mass at *T* = 650 min.

Next, before defining the theoretical model used in the main text to analyse this data, we introduce an angular coordinate better suited to the geometry of the problem. Since differentiation dynamics proceed predominantly along the naso-temporal axis, we define a coordinate along this axis, which we refer to as phase. Mathematically, we define the phase *φ* ∈ [0, 2*π*) as

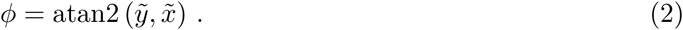

where atan2 is the two-argument arctangent function.

To obtain a coarse-grained description of the differentiation dynamics, we bin the data along the naso-temporal axis into bins of size Δ*φ* = 0.1, resulting in a total of *K* = 63 bins. The resulting matrix of cell counts per phase bin over time serves as the starting point for all subsequent analysis.

### II. MODEL DEFINITION

In this section we introduce a minimal theoretical model to describe the dynamics of a population of *N_T_* cells undergoing cell-autonomous differentiation decisions. Each cell is characterised by a binary state variable

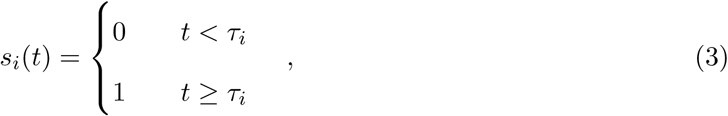

where *t* is the time elapsed since the start of the experiment and *τ_i_* is the intrinsic differentiation time of cell *i*. In our description, cells with *s_i_* = 1 would correspond to Ath5-expressing cells. We assume that, upon activation, Ath5 expression is maintained for the duration of the experiment.

**FIG. 1.**
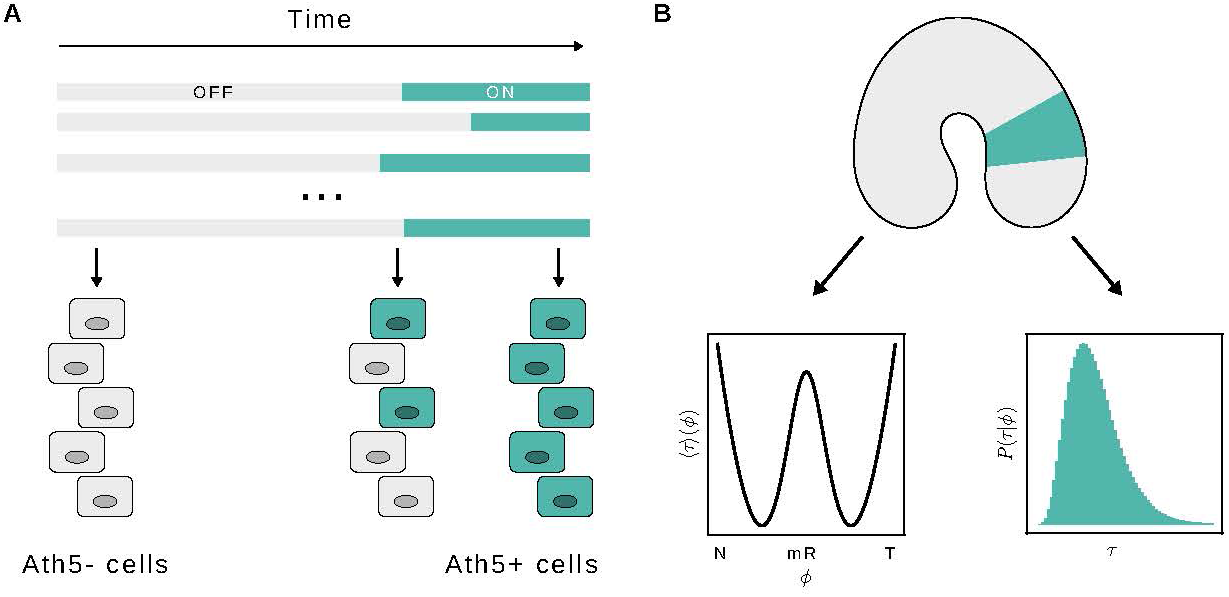
Cartoon depicting the main elements of the model. **A)** The total number of Ath5-positive cells depends on the time elapsed since the start of the experiment and the intrinsic differentiation time of each cell. **B)** Differentiation times *τ* are stochastic variables drawn from a distribution *P* (*τ |φ*) that depends on the naso-temporal coordinate *φ*.

The total number of Ath5-positive cells at time *t* is therefore given by

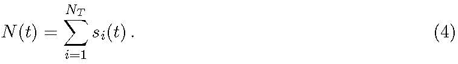

Cell-to-cell variability is incorporated into the model by treating the differentiation times *τ_i_* as stochastic variables drawn from an underlying probability distribution *P* (*τ*). To capture the spatial dependence in the differentiation dynamics, we follow the coarse-graining approach introduced in the previous section and we divide the retina into *K* = 63 regions along the naso-temporal axis, each containing *N_φ_* cells, so that the total number of cells is *N_T_* = *KN_φ_*. Within each region, the differentiation times are assumed to follow the same distribution *P* (*τ |φ*), which depends on the naso-temporal coordinate *φ*. A schematic description of the model is shown in Figure 1.

Since the state of each cell is fully determined by its differentiation time *τ_i_*, drawn from *P* (*τ |φ*), the state *s_i_*(*t*) follows a Bernoulli distribution with parameter *p*(*t, φ*) = *F_φ_*(*t*), where *F_φ_* is the cumulative distribution function (CDF) of the differentiation time distribution *P* (*τ |φ*). Furthermore, since differentiation decisions are cell-autonomous, the total number of Ath5-positive cells within each region is thus distributed as a Binomial distribution of parameters *N_φ_* and *F_φ_*

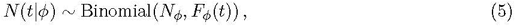

**FIG. 2.**
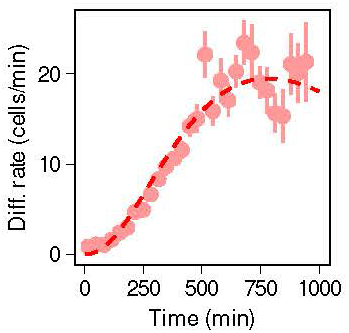
Average differentiation rate *Ṅ* (*t*) across all four control embryos. Error bars represent the standard error of the mean. The red dashed line corresponds to a Gamma distribution fit with parameters estimated via maximum likelihood from the empirical differentiation rate data (*α* = 3.36, *θ* = 331.95 min, *N_T_* = 5163.45).

and the total number of Ath5-positive cells across the whole retina follows a Poisson-Binomial distribution

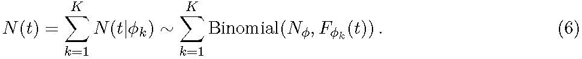

#### A. Differentiation time distribution

Before fitting the model to the experimental data, we need to specify the functional form of the differentiation time distribution. Since the differentiation times completely determine the cell accumulation dynamics, the differentiation rate *Ṅ* (*t*), which measures the number of cells activating Ath5 expression between *t* and *t* + *dt*, provides a direct empirical estimate of the underlying differentiation time distribution in the limit of large cell numbers. We therefore use the empirical differentiation rate across all control embryos as a proxy for this distribution (Figure 2).

We model the differentiation time distribution as a Gamma distribution with shape parameter *α* and scale parameter *θ*. We choose this distribution because the sum of independent Gamma-distributed variables with a common scale parameter is itself Gamma-distributed, which is analytically convenient: it ensures that the distribution of the mean differentiation time across the whole retina retains the same functional form as that of each individual spatial region. Under this model, the empirical differentiation rate is proportional to a Gamma distribution with proportionality constant *N_T_*, the total number of cells. In Figure 2 we show that this approach fits the empirical differentiation rate well. To account for the spatial dependence of differentiation timing, we allow the shape parameter *α_φ_* to vary with the naso-temporal coordinate *φ*, while the scale parameter *θ* is assumed spatially uniform. The differentiation time distribution given the naso-temporal coordinate *φ* then reads

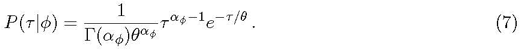

### III. BAYESIAN PARAMETER INFERENCE

Having defined the model and the differentiation time distribution, we estimate the posterior distribution of model parameters given the observed Ath5-positive cell counts along the naso-temporal axis *N* (*t|φ*) using Bayesian inference. We implemented the inference framework in JAGS 4.3.2 using the rjags wrapper [1]. Parameter inference is performed on the first 600 minutes of imaging (120 frames) and the parameter set is given by (*N_φ_, θ,* ***α****_φ_*), where ***α****_φ_* = (*α_φ_*_1_ *,..., α_φK_*).

We place weakly informative priors on all model parameters, centered around values informed by the fit to the aggregate differentiation rate distribution (Figure 2), that respect the positivity of the corresponding parameters. Specifically, we assign a binomial prior to the number of cells per bin *N_φ_* and lognormal priors for the scale and activation time parameters centered near the maximum likelihood estimates 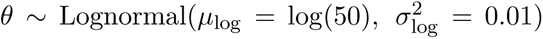 and 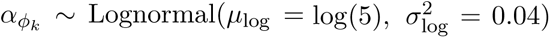. Our results are robust to the precise choice of mean and variance of these priors.

The model was fitted separately to the matrix of cell counts per phase bin over time obtained for each individual eye using four parallel chains, each with an adaptation phase of 5000 iterations followed by a burn-in of 15000 iterations, after which 10000 samples were collected per chain, yielding 40000 posterior samples in total. Figure 3 shows an example of the resulting posterior parameter distributions and pairwise correlations. We provide the full set of distributions for all eyes in a companion compressed file. Across posterior samples ⟨*α*⟩ and *θ* are negatively correlated, indicating that the typical differentiation time across the retina, the mode of the differentiation time distribution *τ ^⋆^* = (*α* − 1)*θ*, is tightly constrained and approximately constant across samples.

Convergence was assessed using the Gelman-Rubin statistic [2] and the multivariate potential scale reduction factor (MPSRF) [3] for the spatially correlated parameters ***α****_φ_*.. The maximum *R̂* across all parameters for each individual eye and the corresponding MPSRF are reported in Table 1.

**FIG. 3.**
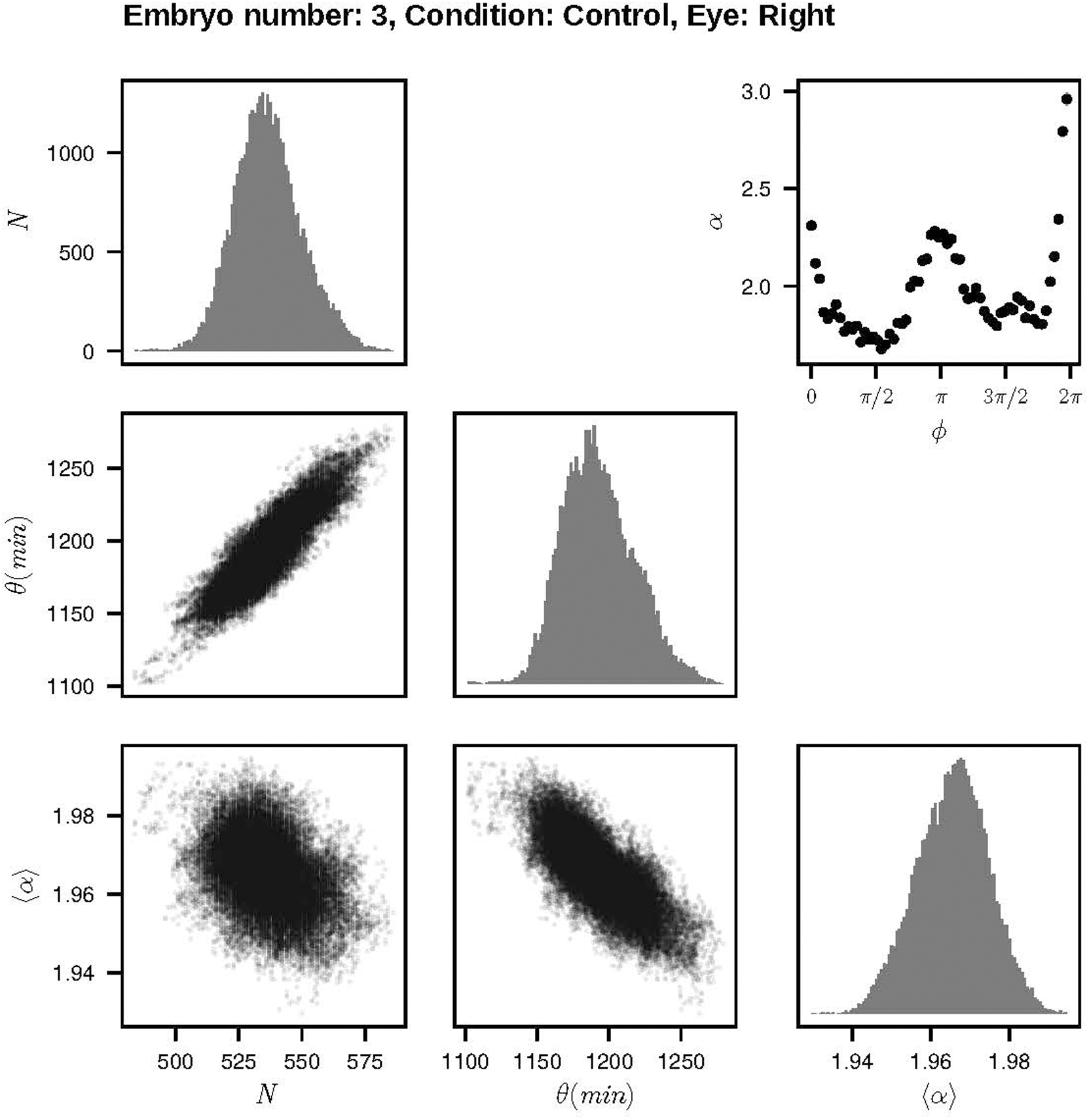
Posterior parameter distributions obtained for the right eye of embryo number 3 of the control condition. Diagonal panels correspond to the marginal distributions of each parameter pooled across chains. Off-diagonal panels show pairwise scatter plots of posterior samples. The upper-right panel shows the mean fitted value of *α* as a function of the naso-temporal coordinate, with error bars indicating the standard deviation of the posterior distribution at each position. Error bars are smaller than the symbols used to the represent the average *α*.

The R implementation of the code used to fit the model can be downloaded from https://github.com/al-sina/bilaterality.

**TABLE I.** Convergence diagnostics obtained from the fitting results for each individual eye.

| Condition | Embryo | Eye | Max $\hat{R}$ | MPSRF |
| --- | --- | --- | --- | --- |
| Control | 1 | Left | 1.083 | 1.039 |
| Control | 1 | Right | 1.093 | 1.066 |
| Control | 2 | Left | 1.033 | 1.014 |
| Control | 2 | Right | 1.059 | 1.043 |
| Control | 3 | Left | 1.019 | 1.018 |
| Control | 3 | Right | 1.125 | 1.070 |
| Control | 4 | Left | 1.063 | 1.040 |
| Control | 4 | Right | 1.014 | 1.010 |
| Ath5-MO | 1 | Left | 1.025 | 1.018 |
| Ath5-MO | 1 | Right | 1.099 | 1.052 |
| Ath5-MO | 2 | Left | 1.160 | 1.095 |
| Ath5-MO | 2 | Right | 1.023 | 1.021 |
| Ath5-MO | 3 | Left | 1.014 | 1.012 |
| Ath5-MO | 3 | Right | 1.045 | 1.020 |
| Ath5/Ptmf1-MO | 1 | Left | 1.064 | 1.044 |
| Ath5/Ptmf1-MO | 1 | Right | 1.045 | 1.032 |
| Ath5/Ptmf1-MO | 2 | Left | 1.058 | 1.042 |
| Ath5/Ptmf1-MO | 2 | Right | 1.057 | 1.042 |
| Ath5/Ptmf1-MO | 3 | Left | 1.043 | 1.032 |
| Ath5/Ptmf1-MO | 3 | Right | 1.060 | 1.043 |

#### A. Assesing the independence of differentiation events from data

To test whether differentiation events observed in the data are independent, we compared the dispersion of empirical differentiation events relative to the dispersion obtained from simulations of the model with gamma distributed differentiation times. We calculated differentiation timings *τ_i_* as the time of emergence of new differentiated cells in the experimental data and, for each differentiation time, we computed the transformed variable

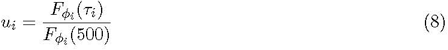

where *F_φi_* is the cumulative distribution function of differentiation times inferred for the spatial region where the differentiation event took place and the denominator is a normalisation factor to account for the size of our observation window.

**TABLE II.** p-values obtained for the observed dispersion statistic for each individual eye of the control embryos.

| Condition | Embryo | Eye | p-value |
| --- | --- | --- | --- |
| Control | 1 | Left | 0.349 |
| Control | 1 | Right | 0.505 |
| Control | 2 | Left | 0.847 |
| Control | 2 | Right | 0.888 |
| Control | 3 | Left | 0.414 |
| Control | 3 | Right | 0.950 |
| Control | 4 | Left | 0.962 |
| Control | 4 | Right | 0.387 |

Assuming that differentiation events are independent draws from a Gamma distribution, the transformed variable *u_i_* is uniformly distributed in [0, 1]. To test whether the empirical differentiation events are consistent with independent differentiation initiation, we binned the transformed values into *M* = 20 equal-width bins and computed the chi-square dispersion statistic

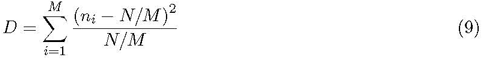

where *n_i_* is the number of events in bin *i*, and compared the observed dispersion statistic against the statistic obtained from simulations of the fitted model to compute empirical p-values for the observed dispersion statistic for each individual eye in our control dataset. We provide the individual p-values obtained from comparing the observed dispersion statistic against 1000 simulations from the fitted model in Table II.

Our results show that, across all eyes, differentiation events showed timings consistent with those expected under the assumption of independent events. Specifically, we find *p >* 0.05 in all cases, providing no evidence against independence.

### IV. RECAPITULATING CONTROL DYNAMICS

The inferred dynamics accurately recapitulate the developmental dynamics of control retinas, as we show in the main text by generating kymographs from stochastic simulations with parameter combinations sampled from the posterior distribution. Moreover, the model can also quantitatively reproduce experimentally measurable quantities such as the average number of Ath5-positive cells and the differentiation rate over time. Taking the average of both sides of Equation 6, the average number of Ath5-positive cells at time *t* reads

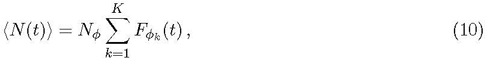

whereas for the differentiation rate we obtain

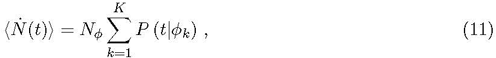

where *P* (*t|φ_k_*) is the differentiation time distribution at the position *φ_k_* along the naso-temporal axis. Model predictions shown in the main text correspond to the analytical expressions derived in this section averaged over 100 parameter combinations sampled from the posterior distribution. To compare with the experimental data in the main text, we align model and experimental results by setting the origin of time at the time point at which the average cell count across the whole retina first reaches 50 cells.

#### A. Predicting intra-embryonic variability

To address whether the observed variability between the two eyes of the same embryo can emerge from the stochasticity of the dynamics, we calculated the squared difference between the number of Ath5-positive cells in the two eyes of the same embryo as a measure of variability. Then, we compared the empirical squared difference against the expected squared difference for two independent realisations of the model described by Equation 6 with parameters obtained from fitting one of the eyes. If we denote by *N*_1_(*t*) and *N*_2_(*t*) the number of Ath5-positive cells at time *t* in two independent realisations the expected squared difference is given by

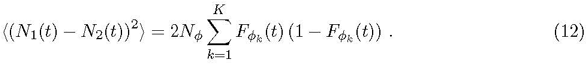

Model predictions shown in the main text correspond to the average over a total of 200 parameter combinations sampled from the posterior distribution, with 100 independent samples per eye.

### V. DIFFERENTIATION TIME PROFILE AND INTRA-EMBRYO VARIABILITY

Motivated by the inferred typical differentiation time profiles of the double morphant embryos, we analyse how the amplitude of the typical differentiation time profile along the naso-temporal axis impacts intra-embryo variability. To this end, we model the spatial dependence of the shape parameter *α_φ_* as a periodic function with minima at 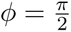 and 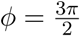 and tunable amplitude

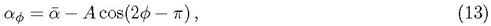

**FIG. 4.**
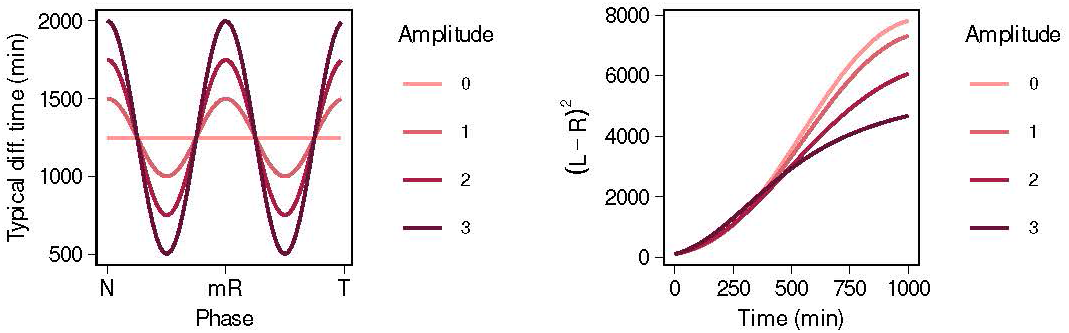
**A)** Typical differentiation time as a function of the naso-temporal coordinate *φ* for different values of the parameter *A* ranging from flat (no spatial dependence) to values larger than the amplitude inferred for control embryos. **B)** Theoretical average squared difference between two independent realisations of the same stochastic process according to Equation 12 for different values of the parameter *A*. Time is normalised such that *t* = 0 corresponds to the time at which the mean Ath5-positive cell count reaches 50. Remaining parameters are fixed to *θ* = 250 min, 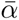 = 6 and *N_φ_* = 250. The number of retinal regions *K* is chosen such that each region has size Δ*φ* = 0.1, as in the processed experimental data.

where 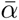 is the mean shape parameter across the retina and the parameter *A >* 0 controls the amplitude of the typical differentiation time profile.

Figure 4 shows that increasing the amplitude of the mean differentiation time profile reduces intra-embryo variability in the early stages of the process. This results from how differentiation is distributed in space and time. A high-amplitude profile staggers differentiation across the retina, so that at any given time only a few spatially localised regions are differentiating. If the differentiation time distribution is not too broad, this limits the number of cells contributing to variability between eyes at that time. A low-amplitude profile, by contrast, makes differentiation timing more uniform across the retina, so a much larger fraction of cells differentiates around the same time, producing a larger peak in variability between eyes.

Over longer time scales, however, Equation 12 predicts a decrease in variability, as both eyes approach the saturation limit of *N_φ_K* Ath5-positive cells (Figure 5). As a result, the initial reduction in variability produced by larger amplitudes compared to flatter profiles, is followed by a later regime in which variability decreases to zero more slowly than in the flatter case.

To quantify the increase in variability in the initial stages caused by the reduced amplitude of the shape parameter ***α****_φ_* profile across the retina in double morphant embryos compared to controls (Figure 5G-H of the main text), we calculated the expected variability between sister eyes given the average scale *θ*_*_ and *N_φ_* parameters across control embryos combined with the average ***α****_φ_* profile from the double morphant embryos.

**FIG. 5.**
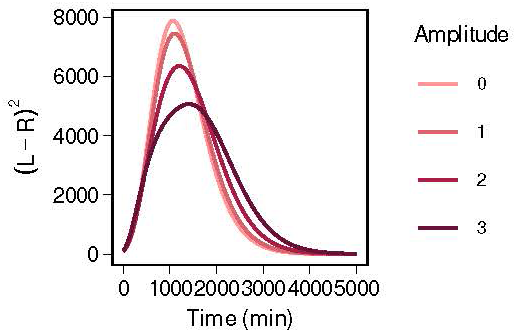
Theoretical average squared difference between two independent realisations of the same stochastic process according to Equation 12 for different values of the parameter *A* over a long time interval. Time is normalised such that *t* = 0 corresponds to the time at which the mean Ath5-positive cell count reaches 50. Remaining parameters are fixed to *θ* = 250 min, 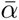 = 6 and *N_φ_* = 250. The number of retinal regions *K* is chosen such that each region has size Δ*φ* = 0.1, as in the processed experimental data.

**FIG. 6.**
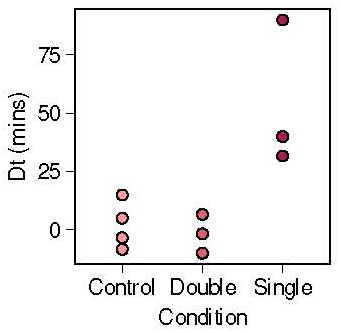
Estimated delay Δ*t* between the right and left eyes of each embryo, calculated according to Equation 14, for the three experimental conditions.

### VI. ESTIMATING INTRA-EMBRYO DELAYS

Since the two eyes of the same embryo are modelled as independent realisations of the same stochastic process, any initial differences between them are maintained over the course of the dynamics. In particular, an initial delay between the two eyes would contribute to the observed intra-embryo variability in the number of Ath5-positive cells.

In the main text, we show that single morphant embryos exhibit an offset between the two eyes. To separate the contribution of this offset from the remaining variability, we estimate the delay Δ*t* between sister eyes as follows. For each eye, we compute the times at which the Ath5-positive cell count reaches the first, second, and third quartiles of its maximum value in log-space. The delay is then estimated as the mean difference between these times across the three quartiles

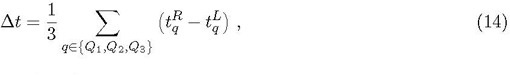

where 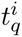 is the time at which eye *i* ∈ *{L, R}* reaches quartile *q*.

In Figure 6 we show the estimated delay for all embryos across the three experimental conditions. Our results show that both control and double morphant embryos small delays of at most a few imaging frames, while single morphant embryos show delays exceeding 30 minutes on average.

